# A Precisely Controlled Long-Acting Immunosuppression Platform Enables Prolonged Survival of Vascularized Composite Allografts

**DOI:** 10.1101/2025.11.07.687240

**Authors:** Sohyung Lee, Fatih Zor, Yalcin Kulahci, John Joseph, Huseyin Karagoz, Shumaim Barooj, Gunjan Malik, Dongsung Park, James N Luo, Lingyun Zhu, Ally Wang, Krisco Cheung, Nancy Li, Purna Shah, Eli Agus, Hayden M Albert, Gopinathan Janarthanan, Sanjairaj Vijayavenkataraman, Priya A Thachet, Helna Mary Baby, Sagi Ravid, Olivia Snapper, Xinyang Chen, Kai Slaughter, Mickael Dang, Ethan Sam, Sam Ho, Amogh Gorantla, Wensheng Zhang, Aadi Kamble, Nutanben Himmatlal Bhingaradiya, Mahima Dewani, Ranit Dutta, Jeffrey M. Karp, Nitin Joshi, Vijay Gorantla

## Abstract

Vascularized composite allotransplantation (VCA) restores complex multi-tissue defects but remains limited by the need for lifelong immunosuppression. Daily oral tacrolimus, the primary maintenance regimen, poses adherence challenges and produces large peak-trough fluctuations within a narrow therapeutic window, resulting in toxicity and periods of under-immunosuppression. Although injectable depots could improve adherence, existing systems frequently generate prolonged supratherapeutic exposure. Most rely primarily on depot matrix properties to govern release kinetics, which alone are insufficient, and in certain systems uncontrolled solvent efflux during phase inversion can exacerbate early burst release. To address these limitations, we developed PRECISE (Programmable, Retrievable, Embedded Controlled ImmunoSuppression Encapsulator), a graft-injectable, in situ-forming depot engineered to precisely regulate systemic tacrolimus exposure. PRECISE integrates a polymeric matrix with rationally selected small-molecule excipients that interact with tacrolimus at the molecular level and modulate solvent dynamics during phase inversion, enabling release control beyond polymer tuning alone. Notably, rapamycin, under select dosing conditions, can also interact with tacrolimus and regulate solvent dynamics, enabling controlled co-delivery of a complementary immunosuppressive combination. This integration of drug-excipient and drug-drug interactions introduces a previously unexplored mechanism for controlling systemic exposure in in situ-forming depots. In a rat hindlimb transplant model, monthly intragraft administration of PRECISE maintained systemic tacrolimus and rapamycin concentrations within the therapeutic window for over 300 days, prolonged graft survival, expanded regulatory T cells, and induced donor-specific hyporesponsiveness. Translational validation in fully MHC-mismatched (SLA Class I and II) pigs, one of the most stringent large-animal transplant models, demonstrated sustained therapeutic drug levels, preserved graft vascular integrity, and graft survival through 90 days. Collectively, these findings establish PRECISE as the first long-acting immunosuppressive platform to maintain systemic drug concentrations precisely within the therapeutic window while enabling controlled dual-drug immunosuppression and the first long-acting immunosuppressive system validated in a fully MHC-mismatched large-animal VCA model, with broad applicability to solid-organ transplantation.

## Introduction

Vascularized composite allotransplantation (VCA) restores complex defects comprising multiple functional tissues such as skin, muscle, bone, and nerves that are critical for mobility and sensation, offering substantial improvement in patient quality of life. Despite surgical success, VCA recipients remain highly susceptible to both acute and chronic rejection due to the immunologic complexity of multi-tissue grafts, with varying antigenicity and vascularization profiles, which complicate immune regulation and heighten the risk of rejection (*1*). Immunosuppressive therapy is therefore essential, with oral tacrolimus (TAC), a calcineurin inhibitor, serving as the mainstay of most VCA protocols.(*2*) However, TAC has a narrow therapeutic window (5-15 ng/mL),(*3, 4*) and post-dose peak concentrations can lead to periods of over-immunosuppression, increasing the risk of nephrotoxicity, malignancy, and opportunistic infection (*4*). Additionally, oral TAC demonstrates substantial interpatient pharmacokinetic variability due to differences in gastrointestinal absorption and first-pass metabolism, resulting in unpredictable drug exposure and alternating periods of under- or over-immunosuppression (*5*). Finally, suboptimal compliance with once- or twice-daily dosing can lead to missed doses and dangerous lapses in immunosuppression, further elevating the risk of graft rejection (*6*). These limitations underscore the necessity for a strategy that can maintain TAC concentrations precisely and stably within the therapeutic window over an extended duration, thereby minimizing the risks of both toxicity and rejection while enhancing patient compliance.

Sustained, long-acting drug delivery strategies represent a promising solution to overcome the limitations of oral immunosuppressive therapies. Formulations administered subcutaneously or intramuscularly can provide prolonged, controlled drug release while bypassing gastrointestinal absorption and first-pass metabolism (*7, 8*). In the setting of VCA, these systems can also be administered directly at or near the graft site, offering the dual opportunity to locally regulate immune activation at the graft interface while maintaining therapeutic systemic drug levels. Graft-embedded or peri-graft controlled-release systems are particularly attractive because they could accomplish both local immunoregulation and systemic immunosuppression (*9*).

Several approaches have been explored for sustained graft-embedded immunosuppression in VCA (*2, 10, 11*). Although these systems have demonstrated preclinical promise in enhancing graft survival and reducing dosing frequency, they fail to consistently maintain systemic TAC levels within the therapeutic window, resulting in periods of supratherapeutic exposure and an increased risk of adverse effects. Most systems exhibit pronounced burst release of TAC and fail to achieve steady-state drug levels rapidly (*12, 13*). This represents a fundamental limitation of many injectable drug delivery systems and is typically addressed by tuning the properties of the bulk polymer matrix. However, this strategy alone is often insufficient to precisely control early-phase drug release. For certain systems, such as *in situ*-forming implants (ISFIs), uncontrolled solvent efflux during depot formation can exacerbate early burst release, as drug partitions into the effluxing solvent and rapid solvent exchange produces highly porous matrices that accelerate drug diffusion (*14, 15*). Together, these challenges define a critical unmet need in VCA immunosuppression: a long-acting injectable platform capable of maintaining systemic TAC concentrations precisely within the narrow therapeutic window.

To address these limitations, we introduce PRECISE – Programmable, Retrievable, Embedded Controlled ImmunoSuppression Encapsulator, a graft-injectable, *in situ*-forming platform designed to overcome burst release and enable precise control of systemic tacrolimus exposure. PRECISE achieves this through an *in situ*-forming poly(lactic-co-glycolic acid) (PLGA) matrix co-formulated with a biocompatible solvent and interactive matrix modulators (IMMs), small-molecule excipients rationally selected to provide a dual function: establishing defined, reversible, non-covalent interactions with TAC while simultaneously modulating solvent efflux. By directly interacting with TAC, IMMs provide molecular-level control over drug retention within the depot, thereby suppressing early burst release and enabling sustained, predictable release kinetics beyond what can be achieved through tuning bulk matrix properties alone. Concurrently, by moderating solvent efflux dynamics, IMMs contribute to controlling the drug release behavior. Upon injection, PRECISE undergoes phase inversion to rapidly form a porous, solid depot at or near the graft site, supporting long-acting, localized immunosuppression while maintaining systemic drug levels within the therapeutic window **(Figure 1)**.

**Figure 1.**
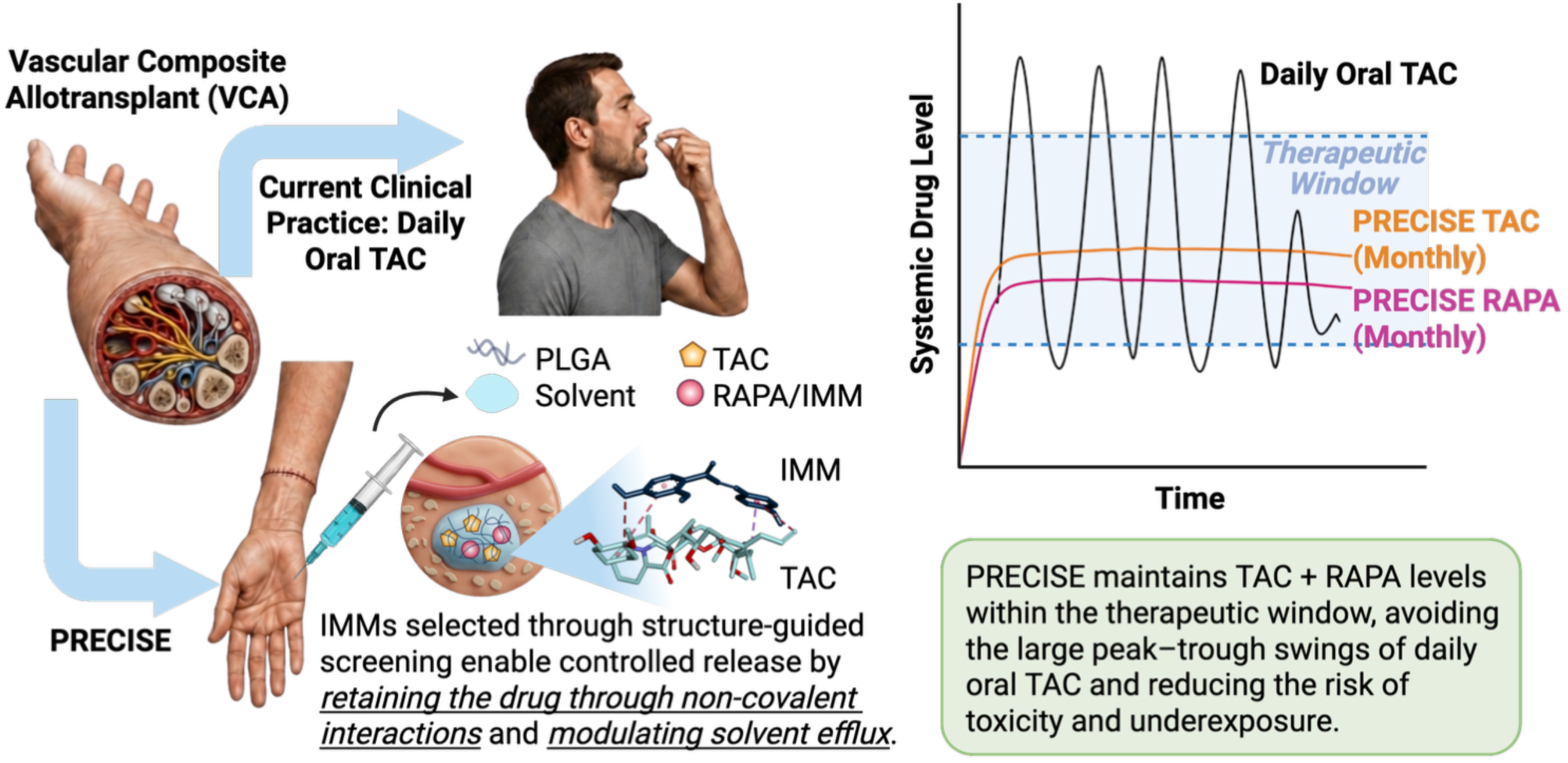
PRECISE - an *In* situ–forming depot for long-term and safe immunosuppression. Oral TAC, the mainstay immunosuppressant in VCA, has a narrow therapeutic window and produces post-dose peaks that increase toxicity risk. Its variable gastrointestinal absorption and first-pass metabolism results in unpredictable systemic levels and alternating periods of under- and over-immunosuppression. In contrast, a single peri-/intra-graft injection of the PRECISE formulation solidifies *in situ* due to solvent efflux to create a localized drug reservoir that results in long-term TAC release with systemic levels precisely controlled within the therapeutic window. IMMs, selected via structure-guided molecular interaction screening are critical to control drug release by interacting with TAC and moderating solvent efflux.

IMMs were identified through a structure-guided screen of FDA’s Generally Recognized as Safe (GRAS) compounds, selected based on predicted non-covalent interactions with key structural features of TAC. GRAS compounds were prioritized for their commercial availability, low cost, and established safety profiles to support scalability and clinical translation. Candidates were first evaluated using *in silico* molecular docking to identify molecules with strong predicted binding affinity for TAC, followed by experimental validation through *in vitro* release studies. Consistent with computational predictions, IMMs exhibiting higher binding affinity, such as epigallocatechin gallate (EGCG), isoliquiritigenin and maltotriose, significantly suppressed initial burst release and reduced overall release rates compared to IMM-free formulations. Importantly, these molecules also attenuated solvent efflux during phase inversion, indicating that drug-excipient binding and solvent modulation act in concert to achieve controlled release. This structure-guided approach provides a prediction-guided basis for excipient selection, with predicted TAC-binding affinity generally tracking the extent of burst suppression observed experimentally. Because IMM selection is based on interaction with the drug rather than solely on modification of the polymer matrix, this design principle may be adaptable to other encapsulated drugs. Together, these findings introduce a mechanistically distinct strategy in which molecular-level drug-excipient interactions are deliberately engineered to regulate release kinetics beyond what can be achieved through bulk matrix tuning alone.

Notably, co-encapsulation of rapamycin (RAPA), another immunosuppressive drug along with TAC at optimized ratios also attenuated TAC release. This effect was mediated by non-covalent interactions between the two macrolides, as well as by a RAPA-dependent reduction in solvent efflux, suggesting that RAPA itself functions as an IMM. This is significant because, in clinical solid-organ transplantation (SOT), mTOR inhibitors such as RAPA or everolimus are often combined with TAC, providing complementary immunosuppressive mechanisms and clinical precedent for combination therapy (*16–18*). These findings demonstrate the formulation flexibility of PRECISE, including single-agent TAC formulations evaluated in vitro and dual-agent TAC+RAPA formulations advanced to in vivo studies, supporting regimen-specific tailoring through structure-guided co-formulation.

PRECISE was well tolerated *in vivo*, with near-complete loss of depot-associated signal by 5–6 weeks after subcutaneous administration. In an orthotopic hindlimb transplantation model, repeated monthly injections of TAC and RAPA co-loaded PRECISE into the graft achieved prolonged, rejection-free graft survival for over 300 days, while maintaining systemic drug levels within the therapeutic window (5-10 ng/mL) and effectively suppressing the initial burst release. Immunological analysis revealed increased circulating regulatory T cells and reduced donor-specific lymphocyte proliferation compared to untreated controls, consistent with durable systemic immunoregulation mediated by sustained, precisely controlled immunosuppression.

To evaluate translational robustness under clinically relevant and immunologically stringent conditions, we tested PRECISE in a porcine heterotopic gracilis flap transplantation model featuring a full swine leukocyte antigen (SLA) Class I and II haplotype mismatch, closely approximating a fully mismatched human VCA. Injection of PRECISE directly into the graft prolonged survival through 90 days, whereas untreated controls exhibited complete rejection by postoperative day (POD) 5. Importantly, systemic tacrolimus and rapamycin concentrations remained consistently within the therapeutic window throughout the study without supratherapeutic excursions. Given the size, vascular complexity, and alloimmune rigor of this large-animal model, these findings provide strong evidence of translational potential and establish PRECISE as a clinically viable platform for achieving durable, precisely controlled long-acting immunosuppression.

Collectively, these findings establish PRECISE as a first-in-class long-acting immunosuppressive platform to maintain systemic drug concentrations precisely within the therapeutic window while enabling controlled dual-drug immunosuppression, and the first long-acting immunosuppressive system validated in a fully MHC-mismatched large-animal VCA model. More broadly, this is the first report to demonstrate that rationally engineered molecular drug-drug and drug-excipient interactions can regulate solvent efflux and program systemic exposure in in situ-forming systems. The underlying design principles may also extend beyond VCA to solid-organ transplantation, where the depot can be placed peri-graft or administered remotely, such as subcutaneously in the abdomen, to achieve long-term immunosuppression with precisely controlled systemic drug levels.

## Results

### Incorporation of Structure-Guided IMMs into PLGA Depots Enables Controlled TAC Release

PLGA-based *in situ* forming depots have been widely explored for long-acting drug delivery and are used in several FDA-approved formulations due to their injectability, biocompatibility, and ability to provide sustained release through polymer degradation (*19, 20*). However, conventional PLGA depots often exhibit an initial burst release phase before reaching steady-state kinetics, which can lead to supratherapeutic drug exposure (*15, 21*). To address this limitation, we hypothesized that incorporating IMMs, small molecules capable of forming non-covalent interactions with TAC, could modulate TAC mobility within the depot, reducing burst release and enabling rapid establishment of a stable release rate. To test this concept, we first performed *in silico* molecular docking studies to evaluate the binding affinity of FDA’s GRAS-listed compounds with TAC. IMM candidates were then incorporated along with PLGA, and their *in vitro* TAC release profiles were compared to formulations lacking IMMs.

To guide IMM selection from the GRAS list, we analyzed the structural features of TAC to identify compounds capable of forming favorable non-covalent interactions. TAC is a highly lipophilic macrolide featuring a 23-membered macrocyclic lactone ring with multiple methyl substituents (e.g., Me 35, Me 39, Me 40, Me 42, Me 43) that make its surface strongly hydrophobic, lowering aqueous solubility. The molecule also contains three isolated C=C bonds (C19=C20, C27=C28, C37=C38) that contribute to its unsaturated, hydrophobic surface, and several oxygenated groups (carbonyl and ether oxygens) that can engage in hydrogen bonding. These features suggested that candidate IMMs with complementary hydrophobic, van der Waals, and hydrogen-bonding interactions would be most effective in forming stable non-covalent TAC-IMM complexes to modulate drug release. Guided by this, we selected eleven GRAS-listed compounds spanning three broad chemical classes **(Figure 2A)**: (i) aromatic polyphenols: EGCG, chalcone, isoliquiritigenin, and 1,3-diphenylpropane, chosen for their π-rich, hydrophobic character; (ii) carbohydrates: β-cyclodextrin, maltotriose, arrowroot starch, and glucose, selected for their ring-based structures, which offer partial conformational compatibility with TAC’s macrocyclic core, and for their abundant hydroxyl groups capable of multivalent hydrogen bonding; and (iii) long-chain hydrophobes: ascorbyl palmitate, octyl gallate, and octylbenzene, selected for their flexible aliphatic tails, which can engage in hydrophobic interactions across TAC’s extensive non-polar surface. We then performed computational docking studies (AutoDock Vina) and analyzed TAC-IMM interaction geometries in PyMOL and BIOVIA Discovery Studio.

**Figure 2.**
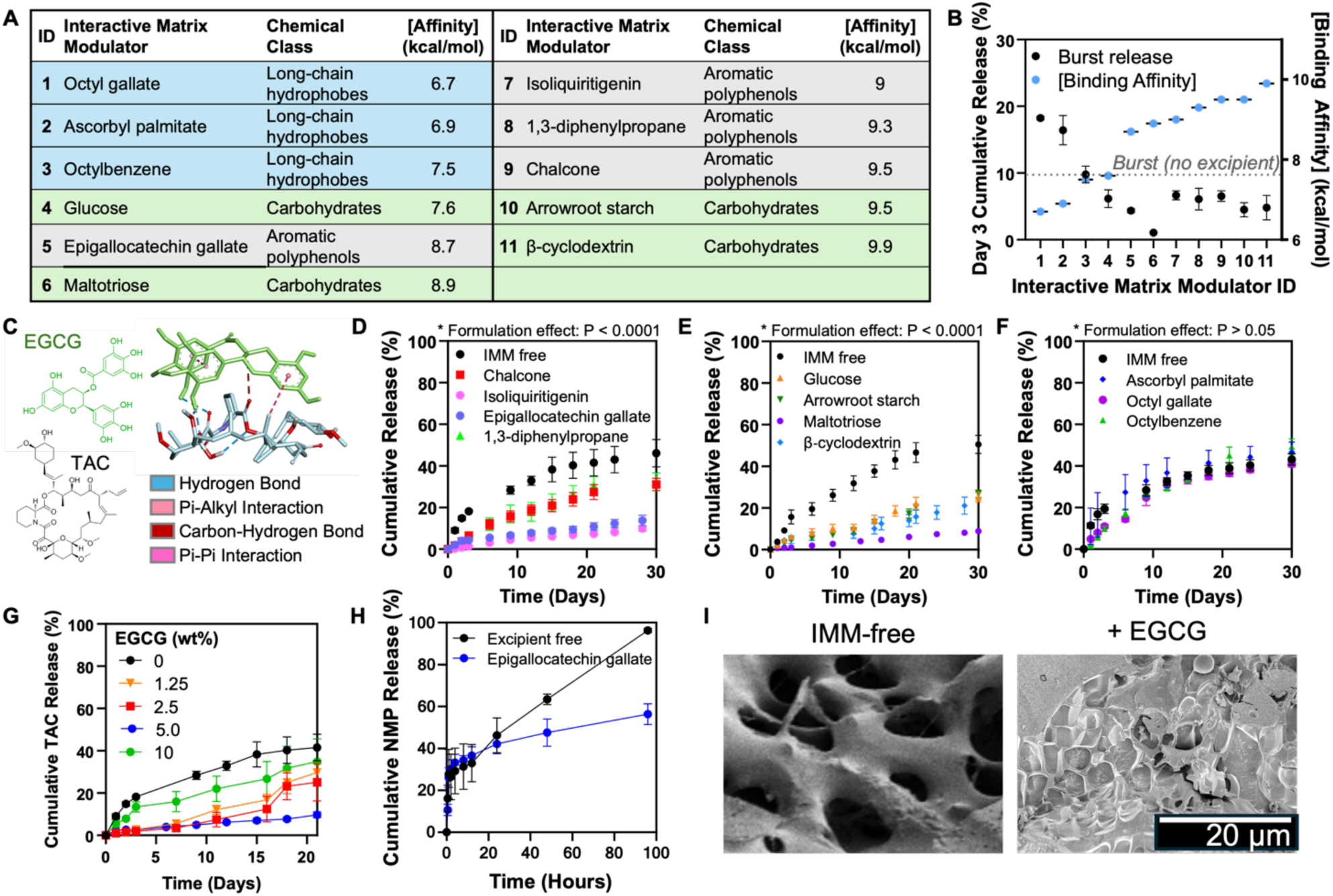
Structure-guided selection of IMMs attenuates TAC burst from PRECISE depots. **(A)** Screen of GRAS-listed IMMs across three chemical classes – aromatic polyphenols, carbohydrates, and long-chain hydrophobes, ranked by predicted TAC-binding affinities from AutoDock Vina (absolute binding-affinity magnitudes are shown; more-negative docking energies indicate stronger predicted binding). **(B)** Plot of Day-3 cumulative TAC release (burst) versus predicted binding affinity for each candidate. Dashed line indicates burst from the no-excipient PRECISE control. **(C)** Representative docking pose of EGCG with TAC. TAC presents hydrogen-bond acceptors and a methyl-rich hydrophobic surface that interacts with EGCG through hydrogen bonding, π–alkyl, and carbon–hydrogen interactions. **(D–F)** *In vitro* TAC release profiles from PRECISE depots doped with 5 wt% IMM, including **(D)** aromatic polyphenols (e.g., EGCG, isoliquiritigenin, chalcone), **(E)** carbohydrates (e.g., maltotriose, β-cyclodextrin, arrowroot starch, glucose), and **(F)** long-chain hydrophobes (octylbenzene, ascorbyl palmitate, octyl gallate). IMM effects were statistically significant for polyphenols and carbohydrates. **(G)** *In vitro* TAC *cumulative* release profiles as a function of EGCG loading (0–10 wt%). **(H)** Cumulative NMP release from PRECISE depots with or without EGCG, showing formulation-dependent NMP efflux. **(I)** Representative cryo-SEM images of TAC-loaded IMM-free PLGA/NMP control and EGCG-containing PRECISE depots at day 7, showing differences in depot microstructure and pore morphology. Cumulative release data in B, D, E, F, G, and H are presented as mean ± standard deviation (n=3). P values in (D–F) were determined using a two-way repeated-measures ANOVA (factors: time and formulation); reported P values correspond to the main effect of formulation. Cryo-SEM images in (I) are representative of n = 3 independent depots per formulation.

Docking analyses revealed distinct TAC-IMM interaction profiles across the eleven candidates **(Figure 2A)**. Binding affinities ranged from –6.7 to –9.9 kcal/mol, with more negative values indicating stronger, more favorable interactions. Complementary *in vitro* release studies were then conducted to experimentally examine whether the interactions between TAC and IMMs could contribute to reducing burst release and extending sustained release relative to the IMM-free formulation. All formulations were prepared using 45 wt% PLGA dissolved in N-methyl-2-pyrrolidone (NMP), a biocompatible solvent, to which 5 wt% of the selected IMM was added relative to the total formulation. TAC was encapsulated at a concentration of 28 mg/mL, selected to deliver a 7 mg TAC dose in a 250 µL injection volume for *in vivo* studies, a dosing regimen that was effective in our previous works (*22*). We particularly paid attention to the cumulative drug release by Day 3 as a parameter representing burst release. IMMs with more favorable (i.e., more negative) binding energies generally reduced the initial burst release compared to the excipient-free formulation **(Figure 2B)**. IMMs exhibiting stronger predicted binding affinity suppressed TAC release more effectively than those with weaker predicted interactions. However, the relationship between burst release and binding affinity was not strictly monotonic, suggesting that additional factors, including other formulation properties, may also influence release kinetics.

Polyphenol-based excipients, including epigallocatechin gallate (EGCG), isoliquiritigenin, and chalcone, reduced burst release and overall release rate, as compared to the IMM-free control **(Figures 2C and D)**, likely due to their aromatic, π-rich structures. EGCG emerged as a representative example **(Figure 2C)**, exhibiting a dual benefit of strong molecular binding (−8.7 kcal/mol) and a greater than 50% reduction in burst release. Molecular docking showed that EGCG donates hydrogen bonds via its phenolic hydroxyls to TAC’s polar atoms (**Figure S1**), while π–alkyl and carbon–hydrogen interactions with methylated regions further stabilized the complex, suggesting dual hydrophilic and hydrophobic engagement.

All carbohydrate-based excipients tested, including maltotriose, β-cyclodextrin, arrowroot starch, and glucose, also suppressed burst release and modestly extended sustained release (**Figures 2B and E)**. Maltotriose, a linear trisaccharide, showed the strongest effect, lowering Day 3 cumulative release to ∼1%, compared to ∼10% in IMM-free control. Docking analysis revealed that maltotriose engages in multiple conventional hydrogen bonds with TAC’s carbonyl and ether oxygen atoms (**Figure S1**), as well as carbon–hydrogen interactions with methyl-rich regions on the TAC surface. One unfavorable acceptor–acceptor contact was also observed but did not appear to disrupt overall affinity, which reached −8.9 kcal/mol.

Excipients with long aliphatic chains had a minimal effect (**Figures 2B and F).** Octylbenzene and ascorbyl palmitate, which exhibited the weakest binding affinities to TAC at −6.7 and −6.9 kcal/mol respectively, formed only limited non-polar contacts. The docking visualizations showed that their interactions were largely confined to van der Waals and nonspecific hydrophobic contacts with TAC’s aliphatic regions, and both compounds slightly increased burst release relative to the excipient-free control (**Figure S1**). This limited interaction profile may explain their poor performance in controlling initial drug diffusion, underscoring the importance of specific binding modes beyond general hydrophobicity.

Next, we evaluated whether the release-suppressing effect of IMMs was dependent on their concentration **(Figure 2G)**. EGCG was selected as a representative IMM for this analysis. Cumulative TAC release decreased progressively with increasing EGCG concentrations up to 5 wt%, indicating enhanced drug retention within the depot. However, further increasing the EGCG content from 5 to 10 wt% led to a partial rebound in TAC release, although overall release remained lower than that of the IMM-free formulation.

In addition to directly interacting with TAC, IMMs also modulated solvent dynamics during depot formation. Both EGCG and β-cyclodextrin reduced cumulative NMP efflux relative to the IMM-free formulation, indicating that IMMs also modulate solvent dynamics during depot formation **(Figure 2H, S3A)**. During the first 24 hours, NMP release from EGCG-containing PRECISE formulations was comparable to that of excipient-free controls. However, beyond this point, EGCG markedly slowed cumulative NMP efflux, with 56.3 ± 4.9% of solvent released by 96 hours compared to 96.3 ± 1.4% in controls. Cryo-SEM analysis further revealed progressive, formulation-dependent differences in depot microstructure during phase inversion. At day 1, no pronounced formulation-dependent differences in pore architecture were evident. Distinct differences emerged by day 3 and became more pronounced by day 7 (**Figure 2I and Figure S4**). The reproducibility of IMM-mediated TAC release suppression was further confirmed across independent in vitro release studies using representative IMM-containing formulations (**Figure S2**).

Formulations incorporating these optimized IMMs were designated as PRECISE depots. The PRECISE platform thus represents an optimized, PLGA-based *in situ*-forming depot that integrates structure-guided excipient selection to achieve precisely controlled, long-acting drug release.

### RAPA Functions as an IMM to Control TAC Release and Enable a Combination Immunosuppression Strategy

Since RAPA, another widely used immunosuppressive agent, possesses a rigid macrolide ring **(Figure 3A)** decorated with multiple methyl groups and isolated double bonds, we hypothesized that it might also function as an effective IMM through non-covalent interactions with TAC. TAC and RAPA are frequently co-administered in transplant medicine (*16–18, 23*), where their complementary mechanisms, calcineurin inhibition by TAC and mTOR pathway suppression by RAPA provide clinical precedent for combination therapy. Accordingly, co-encapsulation of TAC and RAPA within the PRECISE platform was designed to leverage their complementary mechanisms within a single depot, while enabling controlled, localized immunosuppression through RAPA’s additional role as an IMM.

**Figure 3.**
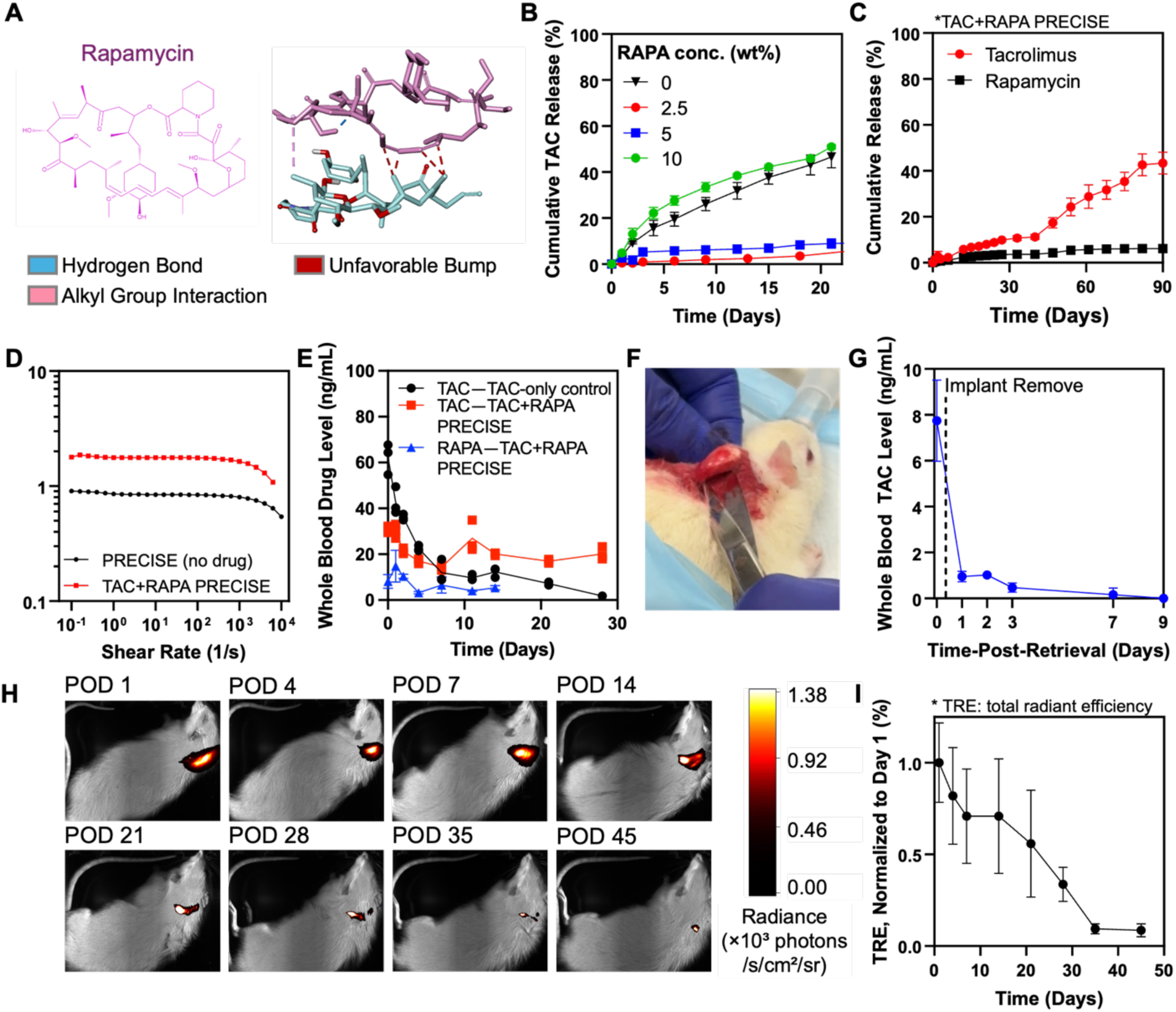
Rapamycin co-encapsulation suppresses tacrolimus burst and enables sustained, retrievable dual-drug delivery from PRECISE depots. **(A)** Representative docking of RAPA with TAC. RAPA, a rigid macrolide bearing methylated alkenes and hydrogen-bond donors, forms hydrogen-bonding and hydrophobic interactions with TAC (predicted ΔG = −12.8 kcal/mol), suggesting stable multimodal complexation. **(B)** Cumulative *in vitro* TAC release from PRECISE with increasing RAPA loading (0, 2.5, 5, 10 wt%). (C) *In vitro* release profile of TAC and RAPA from PRECISE co-loaded with TAC and 2.5 wt% RAPA. **(D)** Representative viscosity data of PRECISE with and without TAC and RAPA (rotational rheometry). **(E)** Whole-blood TAC and RAPA pharmacokinetic profiles in healthy rats following a single subcutaneous injection of TAC-only or TAC+RAPA PRECISE. The assay limit of detection (LOD) for TAC and RAPA was 1.56 ng/mL. For TAC-only PRECISE, TAC was below the LOD in 1 of 3 animals on Day 21 and 2 of 3 animals on Day 28. TAC in the TAC+RAPA group remained detectable through Day 28 and was below the LOD in all animals at subsequent sampling beginning on Day 35. RAPA remained detectable through Day 14 and was below the LOD in all animals by Day 21. **(F)** Photograph showing surgical retrieval of a single PRECISE depot. **(G)** TAC blood levels in rats following PRECISE retrieval, demonstrating controllable drug off-loading. **(H)** Representative IVIS images and normalized total radiant efficiency **(I)** from rats subcutaneously injected with Cy7-labeled PRECISE depots loaded with TAC+RAPA. Data in B, C, E, G, and I are presented as mean ± standard deviation (n=3) Rheology in (D) was performed using n = 3 independent samples per formulation. Representative images in (F) and (H) are from experiments performed in n = 3 rats.

*In silico* docking studies indicated that RAPA’s macrolide ring nests against TAC’s hydrophobic surface and methyl-rich surface **(Figure 3A)**. Simultaneously, RAPA’s hydroxyl and carbonyl groups form conventional hydrogen bonds with TAC’s carbonyl and ether oxygens, yielding a binding affinity of −12.8 kcal/mol. These multimodal contacts suggest that TAC-RAPA pairing could stabilize both molecules within the PRECISE depot.

Consistent with the *in silico* predictions, incorporation of RAPA into the formulation reduced TAC burst release and the overall release rate *in vitro* in a concentration-dependent manner **(Figure 3B)**. At 2.5 wt% RAPA, the Day-3 burst release decreased approximately fivefold, from ∼10% in the RAPA-free control to ∼2%, indicating enhanced stabilization of TAC within the depot. However, at higher RAPA loading (10 wt%), burst release rebounded, surpassing that of the RAPA-free control, a biphasic trend similar to that observed with EGCG. Based on these findings, 2.5 wt% RAPA was selected for subsequent dual-drug PRECISE studies, corresponding to 24 mg RAPA/mL. In this optimized TAC+RAPA co-loaded formulation, both TAC and RAPA were released in a sustained, coordinated manner with substantially reduced early burst release, supporting its use for subsequent in vivo studies **(Figure 3C)**.

The TAC+RAPA PRECISE also exhibited reduced cumulative NMP efflux relative to the RAPA-free formulation **(Figure S3B)**. By 96 hours, 71.4 ± 5.2% of total NMP was released from TAC/RAPA co-loaded depots, as compared to 95.5 ± 0.3 % from RAPA-free controls. Cryo-SEM analysis further showed time-dependent differences in depot microstructure following RAPA incorporation, with differences in pore architecture becoming more pronounced over time (**Figure S4**). This attenuated NMP efflux together with the observed changes in depot microstructure further supports the role of RAPA in modulating depot formation in addition to its interaction with TAC.

Prior to in vivo evaluation, we confirmed the injectability of the optimized TAC+RAPA PRECISE formulation using rotational rheometry across shear rates of 0.1–10⁴ s⁻¹. The viscosity of drug-free PRECISE was 0.97 ± 0.21 Pa·s, which increased to 1.76 ± 0.08 Pa·s upon co-loading with TAC and RAPA. Both values fall within the viscosity range reported for clinically used *in situ* forming depot systems such as Atrigel® (∼1–2 Pa·s) **(Figure 3D)** (*24*), which are compatible with syringe-based administration through 18–25 G needles. These results confirmed that co-loading TAC and RAPA did not compromise the injectability of the formulation.

Having demonstrated the release-modulating effect of RAPA in vitro, we next evaluated whether this effect was retained in vivo by directly comparing TAC pharmacokinetics following a single subcutaneous injection of TAC-only and TAC+RAPA formulations in healthy male Sprague Dawley rats (**Figure 3E**). Both formulations contained the same TAC dose (7 mg per animal), while the TAC+RAPA formulation additionally contained 6 mg RAPA, and whole-blood TAC and RAPA concentrations were monitored through Day 28, with TAC sampling in the TAC+RAPA group extended through Day 56. Consistent with the in vitro release results, the TAC-only formulation showed a greater initial burst, with TAC concentrations of 62.3 ± 6.8 ng/mL compared with 31.0 ± 1.4 ng/mL for the TAC+RAPA formulation at the initial time point. TAC concentrations in the TAC+RAPA group subsequently decreased and remained between approximately 14 and 27 ng/mL from Day 7 through Day 28. In contrast, TAC concentrations in the TAC-only group continued to decline following the initial peak, with concentrations below the assay limit of detection (LOD; 1.56 ng/mL) in 1 of 3 animals on Day 21 and 2 of 3 animals on Day 28. TAC in the TAC+RAPA group remained detectable through Day 28 and was below the LOD in all animals from Day 35 through the final sampling time point on Day 56. This TAC PK comparison further supports the role of RAPA as an IMM in suppressing the initial TAC burst and sustaining TAC release in vivo. Whole-blood RAPA remained detectable through Day 14 and was below the LOD in all animals by Day 21 (**Figure 3E**).

To assess retrievability, we subcutaneously injected TAC+RAPA PRECISE into healthy rats and, after 7 days, surgically removed a single PRECISE depot under brief anesthesia and then monitored TAC levels post-excision. The depot was removed intact, as a single discrete mass, without fragmentation (**Figure 3F**), and circulating TAC declined sharply, falling from 7.75 ± 1.76 ng/mL at the time of excision (Day 0) to 0.95 ± 0.23 ng/mL at 24 h (∼90% decrease). On day 9 following the retrieval, the values approached the assay’s lower quantification limit of 0.16 ng/mL (**Figure 3G**). Together, these data demonstrate that PRECISE offers a controllable, removable depot that could enable rapid clinician-directed cessation or dose reduction in the setting of toxicity, infection, or other clinical needs.

To confirm depot degradation, TAC+RAPA PRECISE depots were formulated using PLGA conjugated to a near-infrared Cy7 fluorophore and tracked *in vivo* by IVIS imaging. The subcutaneously injected depot exhibited a gradual decline in radiance over time, with near-complete signal loss by 5–6 weeks, consistent with substantial degradation of the polymeric matrix **(Figures 3H,I)**. The faster in vivo degradation profile compared with the in vitro drug-release profile likely reflects differences between physiological and in vitro release conditions. This degradation profile and the PK data support the feasibility of monthly to bimonthly dosing intervals for sustained drug delivery.

### PRECISE Enables Sustained Immunosuppression and Long-Term Graft Survival in a Rat Hindlimb Transplant Model

To evaluate the immunosuppressive efficacy of the TAC+RAPA co-loaded PRECISE formulation in preventing graft rejection, we used a rat orthotopic hindlimb allotransplantation model (**Figure 4A**). Transplants were performed from Brown Norway (BN) donors to Lewis recipients, which carry distinct RT1 haplotypes, RT1^n^ and RT1^l^, respectively, that differ across multiple genes within the RT1 complex (*25, 26*). These genetic disparities result in strong incompatibility at the level of antigen presentation, making this model a stringent test of immunosuppressive efficacy.

**Figure 4.**
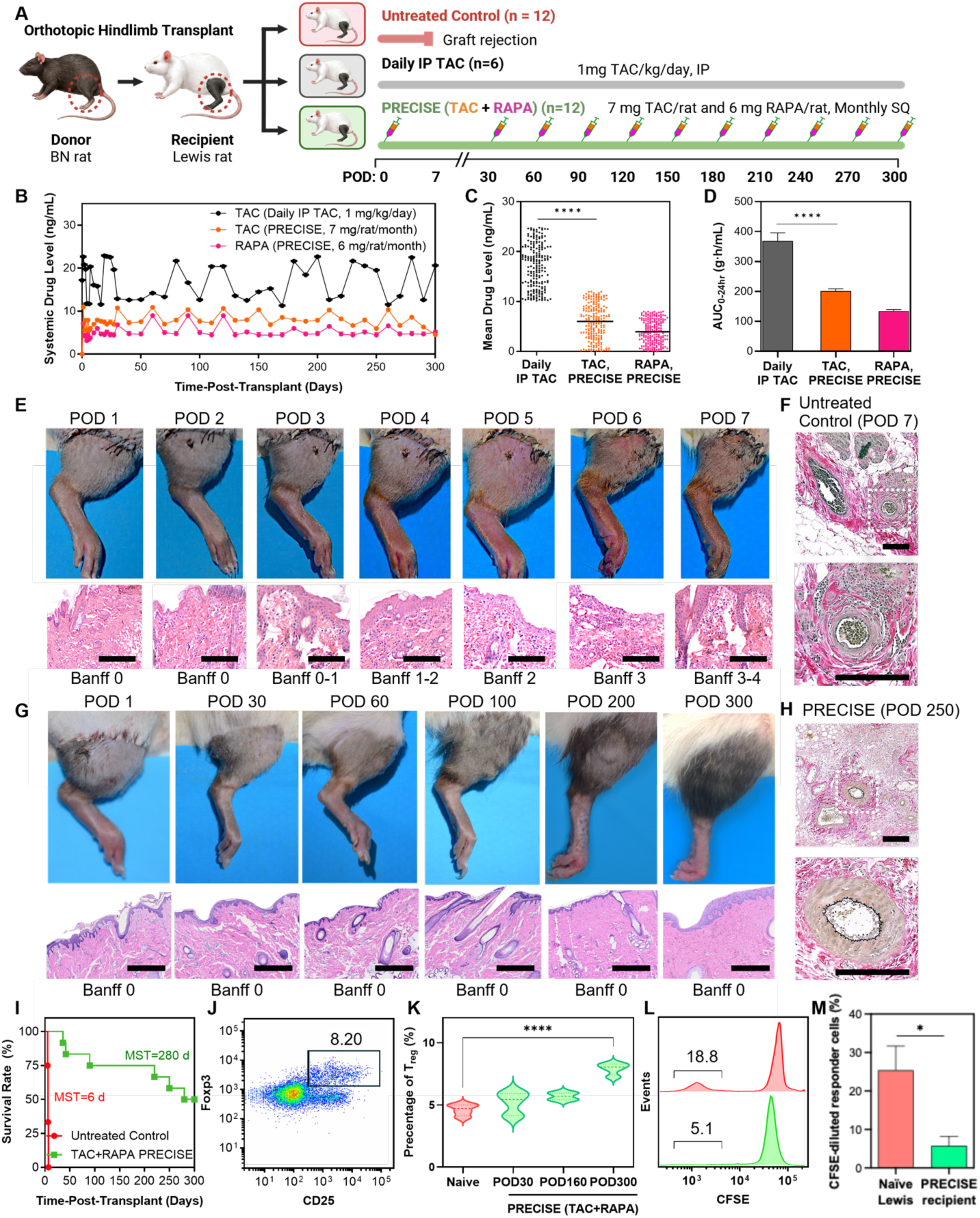
In vivo evaluation of dual-drug PRECISE in a rat limb transplant model. **(A)** Schematic of the study design. Orthotopic hindlimb transplants were performed from Brown Norway (BN) donors to Lewis recipients. Animals received either no treatment (n=12), daily intraperitoneal (IP) TAC (1 mg/kg; n=6), or monthly subcutaneous injections of TAC+RAPA PRECISE (0.25 mL, 7 mg TAC/rat and 6 mg RAPA/rat; n=12). **(B)** Whole-blood drug levels over time in daily IP TAC-treated and PRECISE-treated animals. **(C)** Beeswarm plots showing cumulative whole-blood exposure to TAC and RAPA over the 300-day study period. **(D)** Early systemic exposure measured as AUC0–24 h following a single dose. **(E)** Time-series images showing progressive rejection in untreated recipients, with blackened digits and skin necrosis. H&E-stained tissue sections from untreated controls showing severe acute rejection and necrotic tissue (scale bar: 200 µm). **(F)** Verhoeff–Van Gieson elastin staining in untreated animals on POD 7 showing perivascular infiltration and preserved elastic laminae without intimal thickening (scale bar: 200 µm). **(G)** Time-series images from PRECISE-treated recipients showing graft survival, healing, and full fur regrowth by POD 300. H&E sections from PRECISE-treated animals showing preserved dermal and epidermal architecture at late timepoints (scale bar: 200 µm). **(H)** Elastin staining in PRECISE-treated grafts on POD 250 showing patent vessels with intact elastic laminae and no signs of vasculopathy (scale bar: 200 µm). **(I)** Kaplan–Meier survival curve of untreated and PRECISE-treated recipients post-transplant. **(J)** Representative flow cytometry showing CD4⁺CD25⁺Foxp3⁺ regulatory T cells (Tregs) from a PRECISE-treated animal on POD 300. **(K)** Violin plots showing longitudinal increase in circulating Tregs in PRECISE-treated animals at POD 30, POD 160, and POD 300 compared to naïve controls. **(L)** Representative CFSE-based MLR plots showing donor-specific lymphocyte proliferation in naïve controls and long-term PRECISE-treated recipients after co-culture with irradiated BN stimulator cells. **(M)** Quantification of proliferating CFSE-labeled responder cells across MLR assays (n=3 per group), showing reduced donor-specific lymphocyte proliferation in PRECISE-treated recipients compared with naïve controls, consistent with donor-specific hyporesponsiveness. Data in panels B and D are presented as mean ± standard deviation. Statistical comparisons in C and D were performed using a two-tailed unpaired Student’s t-test. The comparison in M was performed between naïve and PRECISE-treated recipients using a two-tailed unpaired Student’s t-test. The P-value in K was determined using one-way ANOVA.

Three treatment groups were evaluated: (i) untreated controls, (ii) a daily IP TAC comparator group receiving once-daily intraperitoneal (IP) TAC at 1 mg/kg/day, and (iii) the TAC+RAPA co-loaded PRECISE group. In the PRECISE group, animals received a 0.25 mL subcutaneous depot injection at the time of surgery (POD 0), divided between the dorsal and ventral aspects of the transplanted limb. Each depot contained 28 mg/mL TAC and 24 mg/mL RAPA, delivering 7 mg TAC and 6 mg RAPA per animal. Monthly maintenance injections of the same volume and composition were administered through POD 300, for a total of 11 doses per animal.

Because the first dose was applied directly onto the wound bed during surgery, it was essential that the formulation remain localized rather than spreading across surrounding tissue, which would increase exposed surface area and alter release kinetics. This required a shear-thinning system, one that flows readily during injection but rapidly regains viscosity afterward to form a stable depot. To achieve this, 5 wt% palm oil (PO) was added to the standard NMP/PLGA formulation (40 wt% PLGA, 55 wt% NMP, 5 wt% PO), and TAC was pre-encapsulated within a lyophilized triglycerol monostearate (TG-18) hydrogel (80 mg/mL) (*22*) and then incorporated directly into the PLGA phase. Palm oil, a viscous and hydrophobic excipient, was expected to increase overall formulation viscosity and reduce solvent mobility, thereby limiting flow at the injection site. TG-18, a low-molecular-weight amphiphilic gelator, self-assembles through hydrogen bonding and hydrophobic interactions into a weak supramolecular network that behaves as a viscoelastic solid at rest. Under shear, this non-covalent network partially disrupts, leading to a reversible decrease in viscosity that facilitates injection; once shear is removed, the network rapidly reforms, allowing the formulation to re-solidify into a cohesive depot (*27*).

These modifications increased formulation viscosity while imparting desirable shear-thinning behavior **(Figure S5A)**. Because the TG-18 matrix transiently entraps TAC through weak non-covalent interactions, released TAC can still associate with RAPA in the surrounding PLGA phase, preserving the coordinated, controlled release profile characteristic of PRECISE. Consistently, the modified formulation containing palm oil and TG-18 hydrogel exhibited comparable TAC and RAPA release profiles in vitro and pharmacokinetic profiles in vivo to those of the original PRECISE formulation lacking these additives (**Figures S5B–C and S5D–E**). Accordingly, this optimized, shear-thinning formulation was used for all subsequent *in vivo* studies.

In the control group receiving daily IP TAC injections, animals exhibited high and variable blood concentrations throughout the study, with mean levels fluctuating around 19.3 ± 3.8 ng/mL and repeated supratherapeutic peaks exceeding 22 ng/mL at several timepoints (e.g., Days 1, 19, 120, and 200) **(Figures 4B, C)**. These fluctuations highlight the challenge of maintaining stable systemic exposure with bolus dosing. In contrast, in transplant recipients treated with the PRECISE, TAC levels were 10.6 ± 0.64 ng/mL on Day 1, without a discernible post-injection spike, and remained within the 5-10 ng/mL therapeutic range through Day 300 despite repeated monthly injections. RAPA exhibited a similar trend, reaching 7.1 ± 0.61 ng/mL on Day 1 and stabilizing near 5 ng/mL thereafter, supporting the pharmacological relevance of the selected TAC+RAPA PRECISE formulation.

To assess early systemic exposure, we calculated AUC0-24 h for TAC and RAPA using the trapezoidal rule based on plasma concentrations collected from 0 to 24 hours (3-hour intervals) following administration of either PRECISE or daily IP TAC (**Figures 4D, S6**). Early systemic exposure was 368.60 ± 27.11 ng·h/mL for daily IP TAC, 201.11 ± 7.47 ng·h/mL for TAC released from PRECISE, and 134.24 ± 5.22 ng·h/mL for RAPA released from PRECISE. Relative to daily IP TAC, PRECISE reduced TAC AUC0-24 h by more than 40%. These data indicate that the local depot moderates initial systemic exposure while maintaining therapeutic blood levels over an extended period, consistent with the goal of achieving precise and sustained drug delivery with minimized peak fluctuations.

We also evaluated allograft survival to determine the *in vivo* immunosuppressive efficacy of the PRECISE formulation. In untreated controls, all grafts were rejected within 7 days post-transplantation, with a median survival of 6 days **(Figures 4E, I)**. Histopathological analysis of grafts collected at POD 7 revealed severe acute rejection, with Banff grades 3-4 (*28*), extensive tissue necrosis, dense mononuclear infiltration, and vascular injury on hematoxylin and eosin (H&E) staining. Macroscopic examination showed blackened digits and circumferential skin sloughing, consistent with advanced rejection. Verhoeff–Van Gieson (VVG) elastin staining further demonstrated prominent perivascular inflammatory cuffs surrounding vessels with preserved elastic laminae, indicative of perivasculitis without significant intimal thickening **(Figure 4F)**.

In contrast, PRECISE-treated animals exhibited markedly extended graft survival (**Figures 4G, I**). Six of twelve recipients maintained viable grafts throughout the full 300-day study period with no evidence of rejection, and the median survival in the PRECISE group exceeded 250 days, representing more than a forty-fold increase compared to controls. Treated animals showed progressive healing, with recovery of weight bearing and spontaneous use of the transplanted limb during ambulation, together with normal skin coloration, and complete fur regrowth over time (**Figure 4G; Video S1, POD 290**), consistent with long-term graft viability. H&E analysis demonstrated preserved dermal architecture and organized sciatic nerve fascicles with identifiable axon-containing regions in PRECISE-treated grafts. By comparison, the available POD 300 sections from daily IP TAC-treated grafts showed dermal tissue disruption and perineurial/epineurial mononuclear inflammatory infiltrates surrounding sciatic nerve fascicles (**Figure S7**). Consistent with these outcomes, Verhoeff–Van Gieson elastin staining at POD 250 in PRECISE-treated recipients demonstrated patent vascular lumina with continuous elastic laminae, no perivasculitis, and no intimal hyperplasia or graft vasculopathy (**Figure 4H**).

To further assess systemic tolerability during long-term treatment, we analyzed hematology, serum chemistry, and circulating cytokines in the rat VCA study. In daily IP TAC-treated animals, several renal, hepatic, and hematologic parameters changed over time, including elevated creatinine, BUN, ALT, AST, ALP, and total bilirubin, decreased albumin, and progressive anemia and neutropenia. In contrast, these parameters remained within or near reported reference ranges in PRECISE-treated animals across the 300-day study (**Figures 5, S8 and S9)**, and circulating cytokine concentrations likewise stayed within or near baseline ranges (**Figures 5 and S10**).

**Figure 5.**
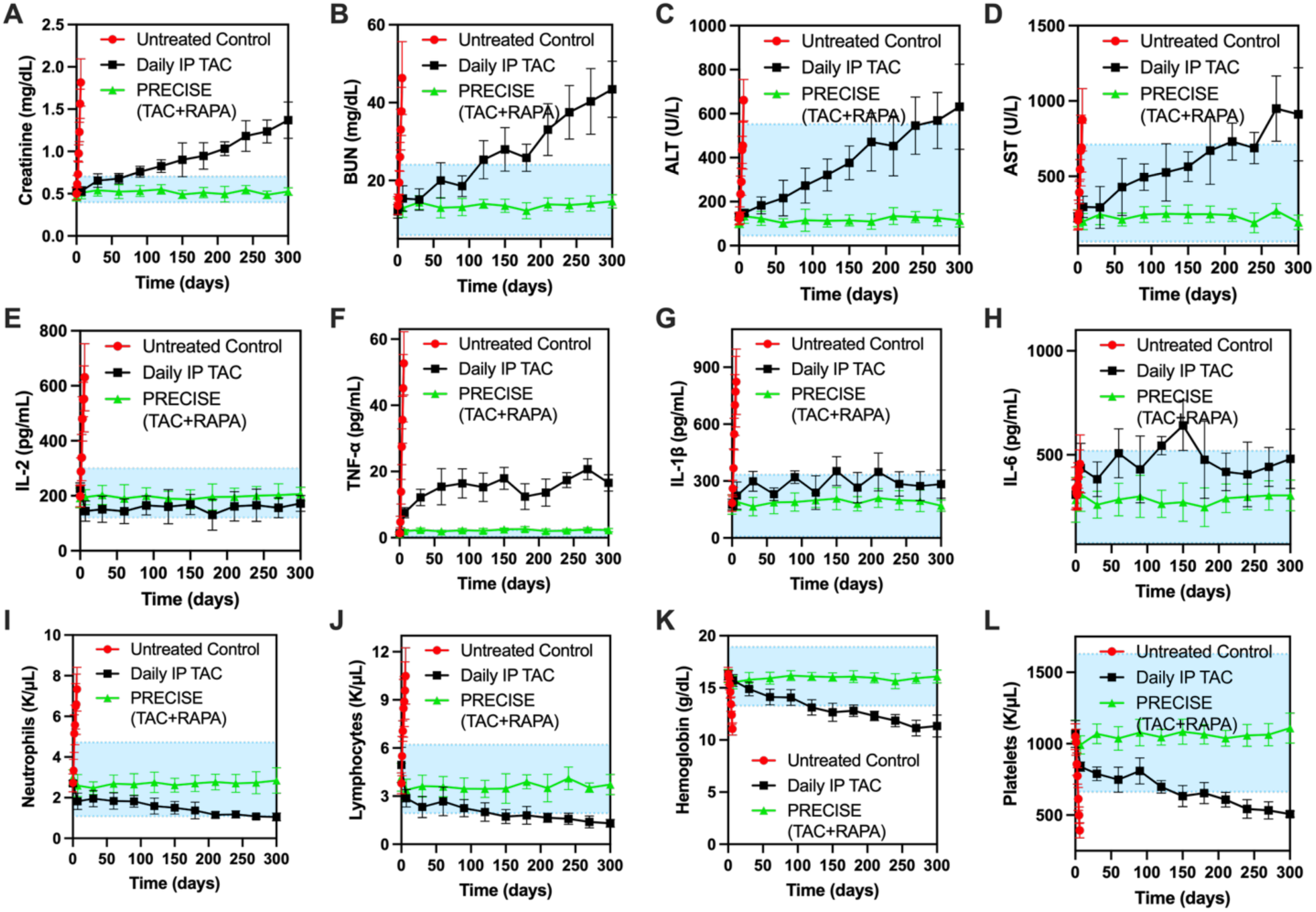
PRECISE maintains serum chemistry, hematologic, and cytokine markers within or near reference ranges during long-term treatment in the rat VCA model. Serum chemistry markers included **(A)** creatinine, **(B)** BUN, **(C)** ALT, and **(D)** AST. CBC parameters included **(E)** neutrophils, **(F)** lymphocytes, **(G)** hemoglobin, and **(H)** platelets. Circulating cytokines included **(I)** IL-2, **(J)** TNF-α, **(K)** IL-1β, and **(L)** IL-6. Daily IP TAC-treated animals showed increases in creatinine, BUN, ALT, and AST and decreases in neutrophils, lymphocytes, hemoglobin, and platelets at later time points, whereas these parameters generally remained within or near reported rat reference ranges in PRECISE-treated animals. Blue shaded regions indicate reported rat reference or baseline ranges where available (*49, 50*). Group sizes were n=12 for untreated controls, n=6 for daily IP TAC-treated animals, and n=12 for PRECISE-treated animals.

To assess immune regulation associated with long-term graft survival, we performed longitudinal flow cytometric analyses of peripheral blood mononuclear cells (PBMCs) in PRECISE-treated recipients versus naïve Lewis rats. Regulatory T cells (Tregs) were defined as CD4⁺CD25⁺Foxp3⁺ populations, using a sequential gating strategy (**Figure S11**). Compared to naïve Lewis rats, PRECISE-treated recipients exhibited elevated frequencies of circulating Tregs at multiple time points post-transplantation. The proportion of CD4⁺ T cells expressing CD25 and Foxp3 increased from a baseline of 4.2 ± 0.4% in naïve animals to 5.9 ± 0.6% at POD 30, 6.3 ± 0.5% at POD 160, and 8.5 ± 0.4% at POD 300 (**Figures 4J, K**). These data indicate an expansion of the Treg compartment in the peripheral blood over time, consistent with a state of active immune regulation in long-term allograft recipients.

To evaluate donor-specific immune reactivity, we performed mixed lymphocyte reaction (MLR) assays using PBMCs isolated from PRECISE-treated recipients at POD 300 and naïve Lewis controls. Responder cells were labeled with carboxyfluorescein succinimidyl ester (CFSE) and co-cultured with irradiated splenocytes from donor BN rats as stimulators. Flow cytometric analysis of CFSE dilution showed markedly reduced lymphocyte proliferation in long-term recipients compared with naïve animals, with 5.8 ± 2.5 % versus 25.4 ± 5.3 % of CFSE⁺ cells undergoing division, respectively **(Figure 4L, M)**. This attenuated proliferative response indicates donor-specific hyporesponsiveness in long-term PRECISE-treated recipients, suggestive of immunologic tolerance or sustained suppression.

Taken together, these findings demonstrate that the PRECISE formulation achieves sustained release of both TAC and RAPA with systemic concentrations maintained within the therapeutic window, resulting in prolonged graft survival and long-term immune regulation characterized by Treg expansion and reduced donor-specific alloreactivity.

### In Vivo Large Animal Evaluation of PRECISE in a Porcine Flap Transplant Model

To assess PRECISE in a clinically relevant large-animal model, we performed allogeneic myocutaneous flap transplantation in miniature swine (mean body weight approximately 25 kg; gracilis free-flap weight approximately 200 g) under conditions of maximal immunological challenge (**Figure 6A**). Donor-recipient pairs were selected based on a compatibility matrix designed to maximize immunologic disparity in their major histocompatibility complex (MHC) antigens while maintaining ABO compatibility (**Figure S12**). Porcine MHC, or the swine leukocyte antigen (SLA), genotyping was performed as previously described (PMID 19392823 and 20121817) with modifications made to the typing primer panels to broaden allele coverage and enhance typing resolution in outbred farm pigs. ABO-incompatible pairs were excluded to prevent hyperacute rejection. Pairs with closely matched SLA were excluded to avoid immunologically permissive transplants. Final selections of donor-recipient pairs prioritized full class I (SLA-1, SLA-2, SLA-3) and class II (DRB1, DQB1) haplotype and antigen mismatches to promote strong alloreactivity. Two reciprocal donor-recipient combinations were selected to control for immunological directionality and to evaluate PRECISE under the most stringent allogeneic conditions.

**Figure 6.**
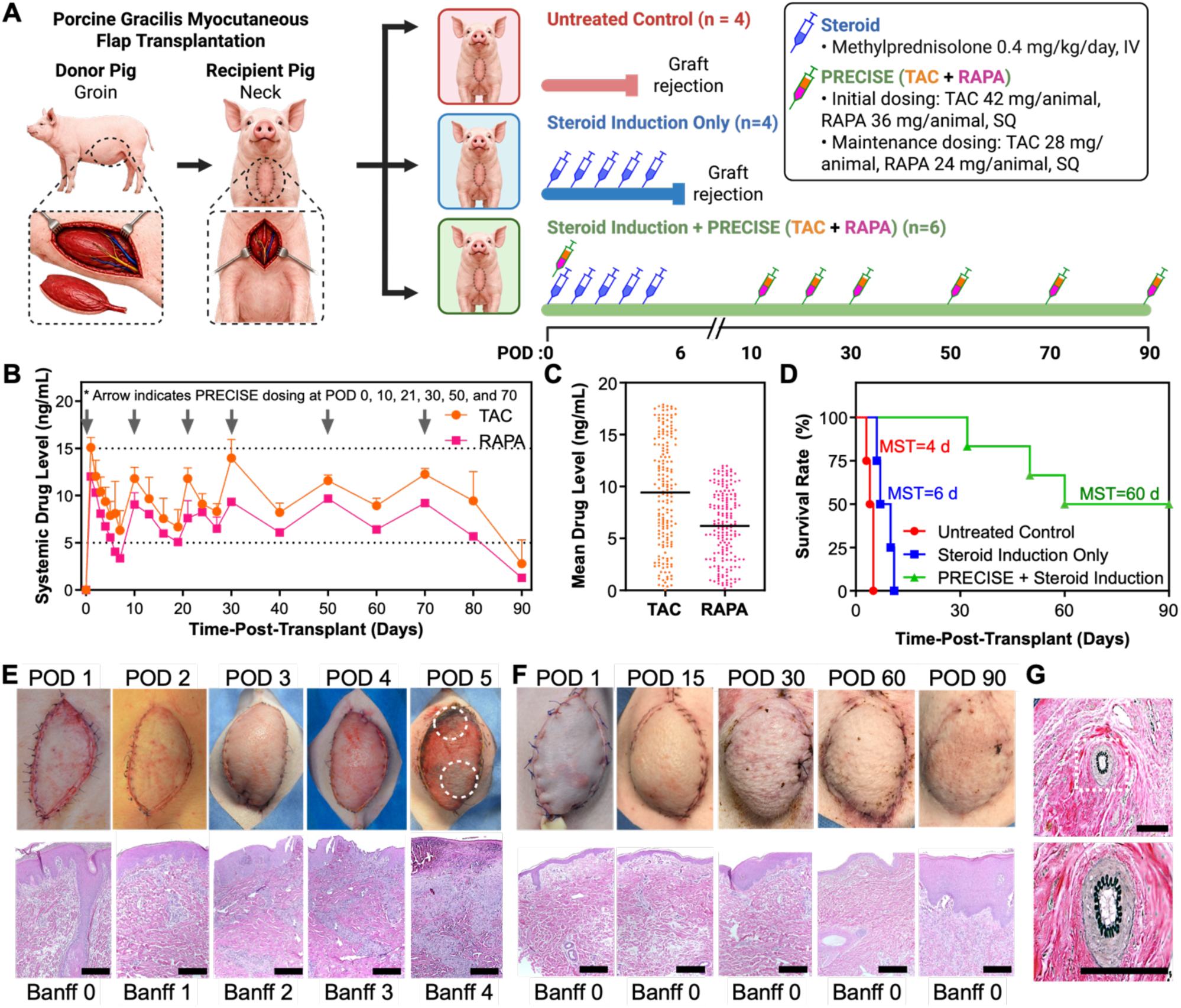
PRECISE enables sustained systemic immunosuppression and prolonged flap survival in a stringent porcine VCA model. **(A)** Schematic of the study design. Myocutaneous gracilis flaps were transplanted between fully mismatched miniature swine donor–recipient pairs. Animals received either no treatment, steroid induction only, or steroid induction plus PRECISE administered at transplantation and on POD 10, 21, 30, 50, and 70, with deposits placed subcutaneously and intramuscularly within the graft. **(B)** Whole-blood drug levels over time in PRECISE–treated animals (black, TAC; magenta, RAPA). **(C)** Bee swarm plots showing cumulative systemic exposure to TAC and RAPA over the 90-day study period in PRECISE-treated animals. **(D)** Kaplan–Meier survival curve of flap grafts. **(E)** Representative gross images and H&E-stained sections from untreated animals at POD 5, showing epidermal necrosis and severe acute rejection (scale bar: 200 µm). **(F)** Gross and histological appearance of PRECISE-treated grafts showing intact architecture and complete fur regrowth through POD 90 with minimal inflammation (scale bar: 45 µm). **(G)** Verhoeff–Van Gieson elastin staining in PRECISE-treated grafts at POD 90 revealing patent vessels with intact elastic laminae and no evidence of vasculopathy (scale bar: 45 µm). Data in B is presented as mean ± standard deviation (sample sizes for each group is indicated in A). Initial group sizes were n = 4 untreated controls, n = 4 steroid-induction-only animals, and n = 6 PRECISE + steroid animals; n at later time points reflects animals remaining in the study.

We evaluated three cohorts under escalating immunosuppressive conditions: (i) untreated control, (ii) systemic steroid induction only, and (iii) TAC+RAPA co-loaded PRECISE plus systemic steroid induction. The steroid induction regimen comprised intraoperative Solu-Medrol (500 mg IV), followed by a once-daily stepwise decrease of 100 mg on postoperative days 1-4, after which steroids were discontinued. PRECISE dosing in the porcine model was determined a priori using a target-tissue scaling framework derived from the rat hindlimb study. The rat formulation achieved therapeutic exposure at a fixed dose of 7 mg TAC and 6 mg RAPA per animal. To translate this to the larger porcine gracilis flap, we normalized by the depot surface-area-to-tissue-mass ratio and applied a conservative downward adjustment to account for slower drug clearance in swine. TAC and RAPA have half-lives of 6–8 hours and 12–20 hours in pigs, respectively, compared with 1–2 hours and ∼6 hours in rats. This scaling approach guided the final induction and maintenance doses. In the PRECISE cohort, PRECISE was administered as follows: at transplantation (induction), six 0.25 mL injections (total 1.5 mL; 28 mg/mL TAC and 24 mg/mL RAPA) delivering a total of 42 mg TAC and 36 mg RAPA per animal (approximately 1.68 mg/kg TAC and 1.44 mg/kg RAPA, assuming a 25 kg animal). Maintenance doses were given on POD 10, 21, 30, 50, and 70, each consisting of four 0.25 mL injections (total 1.0 mL) delivering 28 mg TAC and 24 mg RAPA per animal (approximately 1.12 mg/kg TAC and 0.96 mg/kg RAPA). Injections were distributed across the flap, with three deposits placed subcutaneously and three intramuscularly at the flap corners.

Systemic drug levels were quantified in the PRECISE plus steroid cohort. TAC levels reached 15.08 ± 1.02 ng/mL on POD 1 and were subsequently maintained within 6.33-15.08 ng/mL for the majority of the 90-day observation period (**Figure 6B**). RAPA exhibited an initial concentration of 12.01 ± 0.17 ng/mL on POD 1, followed by sustained levels of 5.57-10.11 ng/mL through most of the study and a gradual decline to 1.29 ± 0.05 ng/mL by POD 90. These data demonstrate that the PRECISE regimen provided prolonged systemic exposure to both immunosuppressants, with concentrations maintained within their respective therapeutic ranges for extended periods (**Figure 6C**).

In the absence of immunosuppression, all grafts failed rapidly, with loss of perfusion at POD 5 (**Figure 6D**). In the steroid-only cohort, all grafts were rejected by POD 11, and survival lasted from POD 6 to POD 11. In the PRECISE plus steroid cohort, all grafts remained viable through POD 32, five of six remained viable at POD 50, four of six at POD 60, and three of six at POD 90. The histopathology corroborated the survival findings. Control grafts sampled at POD 5 demonstrated severe acute rejection (Banff 3–4) with dense mononuclear infiltration, vascular injury, and dermal–epidermal necrosis (**Figure 6E**). In the PRECISE plus steroid cohort, specimens collected at POD 90 retained dermal and epidermal architecture, showed minimal inflammatory infiltrates, and exhibited preserved vascular integrity, consistent with Banff 0–1 (**Figure 6F**). Gross images documented early epidermal necrosis and circumferential skin sloughing in controls and showed progressive healing, restoration of normal skin coloration, and complete hair regrowth in the PRECISE–treated cohort. To further assess vascular integrity, Verhoeff–Van Gieson elastin staining in treated recipients at POD 90 revealed intact elastic laminae, open lumens, and an absence of perivascular inflammation or vasculopathy, indicating preserved vascular architecture without evidence of chronic rejection (**Figure 6G**).

Systemic tolerability was further assessed by hematology, serum chemistry, and multiplex cytokine profiling in the porcine VCA study. CBC parameters and renal and hepatic chemistry markers— creatinine, BUN, ALT, AST, ALP, GGT, and total bilirubin—remained within or near reported porcine reference ranges throughout the 90-day study in PRECISE-treated animals (**Figures 7, S13 and S14**), and cytokine profiling did not indicate sustained systemic inflammatory activation (**Figures 7 and S15**). Untreated and steroid-only grafts, by contrast, showed early increases in renal chemistry markers before rejection on POD 5–11.

**Figure 7.**
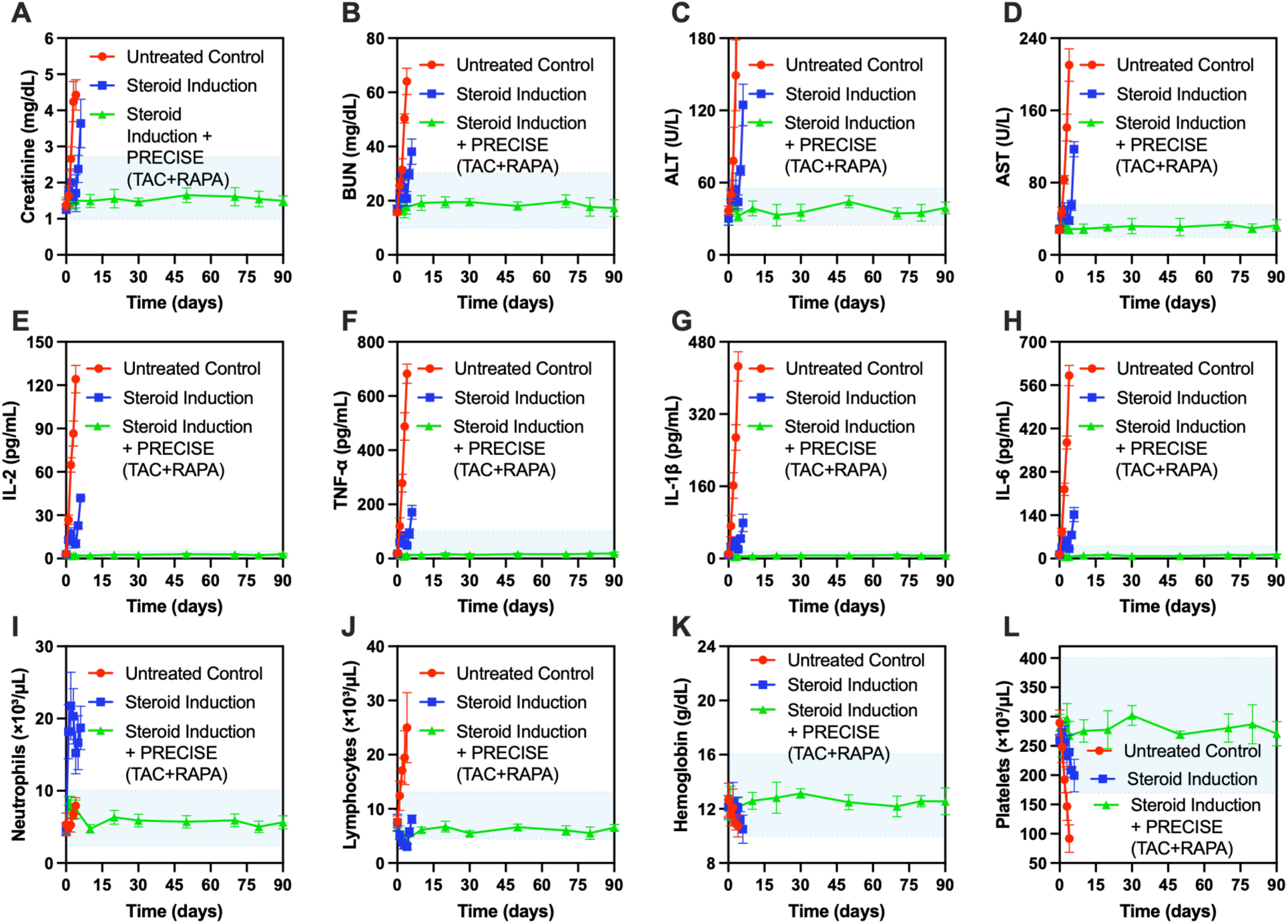
PRECISE maintains serum chemistry, hematologic, and cytokine markers within or near reference ranges in the porcine VCA model. Serum chemistry markers included **(A)** creatinine, **(B)** BUN, **(C)** ALT, and **(D)** AST. CBC parameters included **(E)** neutrophils, **(F)** lymphocytes, **(G)** hemoglobin, and **(H)** platelets. Circulating cytokines included **(I)** IL-2, **(J)** TNF-α, **(K)** IL-1β, and **(L)** IL-6. These parameters generally remained within or near reported porcine reference or baseline ranges in PRECISE-treated animals throughout the 90-day study. Blue shaded regions indicate reported porcine reference or baseline ranges where available (*51*). Group sizes were n=4 for untreated controls, n=4 for steroid-induction-only animals, and n=6 for PRECISE + steroid-treated animals.

Flow cytometry of PBMCs in the porcine PRECISE + steroid cohort showed a progressive rise in circulating regulatory T cells (CD4⁺CD25⁺Foxp3⁺), increasing from ∼5% at POD 0 to ∼11% by POD 80 (**Figure S16**). To enable terminal tissue collection while minimizing repeated invasive sampling, we used a cross-sectional, serial-sacrifice design with one animal per time point, consistent with the principles of Replacement, Reduction, and Refinement (3Rs) and the ARRIVE 2.0 guidelines. Accordingly, these data are descriptive and not powered for statistical (longitudinal) inference; confirmation in larger longitudinal cohorts is needed. These descriptive data are directionally consistent with the rat results and with the extended graft survival observed in this study.

## Discussion

VCA remains constrained by toxicities and the variations in systemic levels of orally administered calcineurin-inhibitor-based regimens such as TAC. TAC has a narrow therapeutic window (∼5-15 ng/mL), exhibits high inter- and intrapatient pharmacokinetic variability from absorption and CYP3A metabolism,(*29*) and therefore requires intensive therapeutic drug monitoring; excursions above and below target are linked to nephrotoxicity or infection risk and rejection, respectively (*5, 30, 31*). In this study, we present PRECISE, a graft-embedded, long-acting formulation that enables precise and sustained immunosuppression through structure-guided control of TAC release. Although excipient-based strategies have previously been used to reduce burst release from PLGA in situ-forming systems,(*15*) these approaches have generally focused on empirically modifying formulation or matrix properties. PRECISE instead uses in silico drug–excipient interactions to guide IMM selection before formulation, with predicted binding affinity generally corresponding to the extent of TAC burst suppression observed experimentally. RAPA also functions as an IMM, and when co-formulated with TAC at select concentrations, it not only regulates TAC release kinetics but also provides complementary mTOR-mediated immunomodulation, resulting in durable graft protection. The *in situ* forming injectable depot maintained stable systemic TAC and RAPA levels and prolonged graft survival in VCA models, through 300 days in rats and 90 days in pigs, without clinical or histologic evidence of rejection.

Various drug-delivery systems have been developed as carriers for long-acting immunosuppressive therapy in VCA, ranging from solvent-based intragraft drug solutions (*11*) and hydrogels (*10, 12, 22*) to microspheres/nanoparticles (*32–35*) and preformed biodegradable TAC-releasing implants (*36, 37*). PRECISE offers several key advantages over these earlier approaches. It substantially reduces burst release and establishes stable, steady-state blood levels shortly after administration. Many prior systems on the other hand, exhibit substantial burst release and limited control over long-term kinetics (*12, 13, 38*). For example, a triglycerol monostearate (TG-18) hydrogel previously developed by our group achieved long-term TAC release and extended VCA survival beyond 100 days in rats following a single injection. (*22*) However, this system produced an initial mean blood TAC concentration of ∼127 ng/mL, an order of magnitude higher than the ∼10 ng/mL observed with PRECISE, before gradually declining toward therapeutic levels. In the porcine model, TG-18 similarly extended graft survival to 90 days but exhibited an early peak of ∼40 ng/mL, whereas PRECISE maintained an initial concentration near 15 ng/mL and achieved near-steady-state plasma levels from day 1. In contrast, the TG-18 hydrogel required approximately 20 days to approach steady-state concentrations in rats and failed to achieve steady-state levels in pigs.

Another group reported a single injection of PLGA microspheres co-encapsulating TAC with mycophenolate mofetil and prednisone, which prolonged median graft survival to >150 days in a rat hindlimb transplantation model (*39*). However, this formulation produced an excessive initial burst for all three drugs, remained supratherapeutic for most of the release period, and failed to achieve stable steady-state plasma concentrations. In contrast, PRECISE rapidly achieves and maintains stable therapeutic concentrations of both TAC and RAPA while substantially reducing burst release, underscoring its distinct pharmacokinetic precision. To our knowledge, this represents the first report of a dual-immunosuppressive, long-acting drug-delivery platform that maintains systemic drug concentrations precisely within the therapeutic window.

Furthermore, PRECISE is retrievable through a small skin incision. Upon injection, the formulation undergoes phase inversion from liquid to solid, resulting in a localized implant that can be surgically excised if rapid drug cessation is required. By contrast, solvent-based intragraft solutions (e.g., DMSO-based) (*11, 40*) and dispersed microspheres/nanoparticles (*32–35*) are not practically retrievable once injected, and many tissue-infiltrating injectable hydrogels (*10, 12, 22*) likewise cannot be fully removed. Given TAC’s risk of toxicity and infection, this capability for rapid discontinuation represents a clinically meaningful safety advantage of PRECISE.

PRECISE achieved rapid steady-state drug levels and substantially reduced burst release both in vitro and in vivo through structure-guided co-formulation of TAC with small-molecule IMMs. Although excipients have previously been used to mitigate burst release from PLGA in situ-forming systems, these approaches have largely focused on modifying formulation or bulk matrix properties (*15*). Injectable PLGA depots also undergo solvent exchange and phase inversion after administration, during which rapid solvent efflux and pore formation can contribute to early burst release (*14, 15*). The IMMs formed non-covalent interactions with TAC that likely reduced its diffusivity within the polymer matrix, while their presence also modulated solvent efflux during phase inversion. The marked reduction in early TAC release was observed even when cumulative NMP release during the first 24 hours was similar between EGCG-containing and IMM-free formulations, supporting a substantial contribution from direct TAC–IMM interactions to the observed release control. Cryo-SEM analysis further showed progressive formulation-dependent differences in depot microstructure for both EGCG- and RAPA-containing PRECISE, with differences in pore architecture becoming more pronounced during depot formation. Previous studies of PLGA in situ-forming systems have similarly shown that solvent exchange and phase-inversion kinetics can influence the resulting pore structure and drug release (*14, 41–43*). The formulation-dependent differences in NMP efflux may reflect changes in solvent/non-solvent exchange during phase inversion, which can be influenced by formulation composition and can alter the resulting depot morphology, porosity, and drug release behavior (*14, 15, 41–43*). Together, these findings support a combined mechanism in which direct TAC–IMM interactions and formulation-dependent changes in depot formation contribute to control of TAC release. The effect of IMMs was concentration-dependent; for both EGCG and RAPA, we observed reduced burst release and slower overall release kinetics at 2.5-5 wt%, whereas the effect was lost at 10 wt%. We hypothesize that this non-monotonic behavior arises from excipient saturation within the polymer matrix leading to the disruption of the depot microstructure at higher IMM loadings.

Importantly, translational robustness of PRECISE was demonstrated in a fully MHC-mismatched (SLA Class I and II) porcine gracilis flap transplantation model, which closely recapitulates the surgical complexity, tissue composition, and immunologic stringency of clinical VCA (*44*). Large-animal VCA models are considerably more demanding than rodent systems, with greater graft mass, vascular load, and pharmacokinetic variability. The ability of PRECISE to maintain therapeutic TAC and RAPA concentrations within the target window from day 1, preserve graft vascular integrity, and support graft survival through 90 days under these stringent conditions underscores its translational potential. To our knowledge, this represents the first evaluation of a long-acting immunosuppressive platform in a fully MHC-mismatched large-animal VCA model. In addition to evaluating efficacy, we assessed tolerability in both models. Although PLGA/NMP in situ-forming depots have clinical precedent,(*20, 45, 46*) graft-local administration into transplanted tissue represents a distinct setting from conventional subcutaneous depot delivery, and we therefore evaluated the formulation directly in this setting. PRECISE-treated animals maintained hematologic, renal, hepatic, and inflammatory markers within or near reference ranges across both models, supporting local and systemic tolerability alongside the efficacy described above. Functional recovery was also supported by spontaneous weight bearing and use of the transplanted limb during ambulation, with prospective functional scoring in future studies providing a quantitative assessment of motor recovery.

Despite these encouraging results, additional studies are needed to address remaining questions. First, the dosing intervals used here were intentionally conservative to prioritize graft stability rather than to define the maximal duration achievable with a single depot. In our models, TAC and RAPA depots were administered approximately once per month in rats and every 2-4 weeks in pigs to maximize graft success. Even at these intervals, dosing every two weeks or once per month represents a substantial improvement over the current daily oral regimen. Less frequent dosing may be feasible; future studies in healthy large animals should therefore quantify the duration of exposure following a single injection and establish redosing criteria based on pharmacokinetic monitoring and early clinical or histologic indicators of rejection. If needed, formulation parameters such as higher molecular weight PLGA, modified lactic:glycolic ratios, or increased depot volume can be optimized to further extend dosing intervals while maintaining precise systemic control.

Second, additional studies on single-drug PRECISE depots co-formulated with IMMs would be valuable. We observed that RAPA itself functioned as an IMM, suppressing TAC burst release and promoting controlled sustained delivery of both drugs. Owing to this intrinsic effect, subsequent studies primarily focused on the optimized TAC+RAPA PRECISE formulation, including the in vivo efficacy studies. Consequently, we did not perform detailed single-agent evaluations of TAC-only depots co-formulated with small-molecule IMMs such as EGCG or maltotriose, despite their excellent *in vitro* performance. Because many patients are maintained on TAC monotherapy in clinical practice, future work should investigate single-drug PRECISE depots with IMMs to better define *in vivo* dose-response relationships, release durability, safety, and the contribution of depot-mediated TAC delivery without the added contribution of RAPA. Similarly, for combination therapy, the TAC:RAPA ratio can be further tuned as the formulation is adapted for specific transplant indications and dosing strategies, rather than being fixed by the ratio used in this proof-of-concept formulation.

Third, while our porcine model offers clinically relevant skin and soft-tissue architecture under stringent histocompatibility mismatch, the current study was not designed to fully optimize the clinical immunosuppressive regimen or isolate the relative contributions of local graft exposure and systemic drug exposure. We did not include a matched systemic TAC+RAPA comparator or a remote-site PRECISE depot group, both of which would help define whether graft-local delivery provides benefits beyond maintaining therapeutic systemic drug levels. Because systemic combination dosing, route, and target exposure are not standardized in porcine VCA, these comparisons will require dedicated dose-matched studies. This challenge is particularly relevant in outbred, fully mismatched pigs, in which immunosuppressive regimens at doses comparable to those used clinically in humans may be insufficient and more intensive combination therapy can be required (*47*). Future work should therefore compare graft-local PRECISE with remote depot placement and optimized systemic TAC+RAPA regimens, while measuring blood drug concentrations, graft tissue drug levels, immune responses, histopathology, and systemic toxicity. Additional studies in non-human primate (NHP) VCA models (for example, rhesus or cynomolgus macaques) could further align with human drug metabolism and therapeutic drug monitoring, and enable evaluation of steroid-sparing regimens, retrieval protocols, and long-horizon safety. These studies would help refine dose scaling, depot placement, and redosing criteria ahead of early-phase clinical trials.

In summary, PRECISE establishes a simple yet mechanistically engineered platform for achieving controlled, sustained, and clinically actionable immunosuppression in VCA. Through structure-guided coupling of TAC with IMMS, PRECISE substantially reduces burst release, rapidly achieves stable therapeutic concentrations, including for dual-drug immunosuppression, and sustains long-term graft survival without daily systemic therapy. The depot is fully retrievable, providing an additional layer of safety and clinical control for translation. To our knowledge, this is the first demonstration of a long-acting combination immunosuppressant depot capable of maintaining systemic exposure within the therapeutic window while providing durable graft protection. Beyond VCA, this platform may be broadly applicable to solid-organ transplantation, where precise and sustained immunosuppression remains a central challenge.

## Materials and Methods

### Animals

Male Sprague Dawley rats (200–250 g; 8–10 weeks old) were used for healthy-rat pharmacokinetic studies. Male Lewis rats (200–250 g; 8–10 weeks old) were used as recipients for rat VCA studies. Brown Norway (BN) and Fischer 344 (F344) rats of similar age and weight served as donors for acute- and chronic-rejection models, respectively. Animals were maintained under controlled temperature, humidity, and a 12-hour light/dark cycle with ad libitum access to food and water. All procedures were approved by the IACUC and the Department of Defense ACURO and conformed to the NIH Guide for the Care and Use of Laboratory Animals.

Large-animal studies were performed in domestic crossbred swine (Sus scrofa domesticus; Landrace × Yorkshire × Duroc, “L × Y × D”; 6–8 weeks old; mean body weight approximately 25 kg) to capture natural outbred heterozygosity within the SLA complex. All porcine procedures were approved by the institutional IACUC and ACURO and performed under veterinary supervision in compliance with the Guide for the Care and Use of Laboratory Animals (8th edition).

### Preparation of PRECISE

The pre-polymer solution was composed of 40 w/v % PLGA (L:G 85:15, 10-15 kDa, Poly Scientific) and TAC at a concentration of 28 mg/mL. Depending on the presence of excipients, the formulation contained either 55 v/v % NMP (Sigma-Aldrich) with 5 w/v % excipients or 60 v/v % NMP without excipients. All components were added to an 8 mL glass vial and heated until completely dissolved. The formulation was then drawn into a syringe and injected through 23G needle either into a sink consisting of 20% methanol in PBS for in vitro studies or subcutaneously into the transplantation site in animals for in vivo studies. Following administration, formulation underwent solvent exchange, polymerizing into a solid depot.

### In vitro drug release

100 uL of drug-loaded pre-polymer mixture (40w/w% PLGA, 55% or 60%w/w% NMP, and 28mg/ml TAC) was injected into the bottom of a 50mL Falcon tube using a syringe fitted with a 23G needle. Then the sink medium was added. For TAC-loaded PRECISE, we used 20% methanol in water as the sink medium due to low water solubility of the drug. The Falcon tube containing PRECISE depot was then incubated at 37 °C in a shaker incubator. The release medium was collected at pre- determined time points to be analyzed on HPLC.

The collected release sample was subjected to 24h freeze-drying to remove solvents and then reconstituted in 1ml ACN (Sigma-Aldrich). The drug concentration was determined using HPLC (Agilent 1260 Infinity II, USA) with a ZORBAX 300SB-C18 column (Agilent, 3.0 x 150 mm, 3.5 µm) at 40 °C. Chromatographic separation was achieved by gradient elution method, as described in **Table S1**. Calibration standards were prepared in the range of 0.78 - 400 mg/ml and cumulative drug release was calculated accordingly.

### Injectability characteristics

A rheometer equipped with a parallel plate with a diameter of 20 mm and a gap of 1 mm was used to determine the dynamic viscosity of the pre-polymer at room temperature. The solutions were pipetted onto the rheometer, and any excess solution was removed before measurement. The viscosity of the formulation was measured as the shear rate was swept from 1 to 1000 s^-1^. The force required to inject 1 mL of ISFI pre-polymer solution in 30 seconds was calculated using the Hagen-Pouiselle equation:

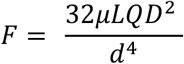

Where μ is the dynamic viscosity of ISFI formulation determined from the rheometer, L is length of needle (15 mm), Q is volumetric flow rate (2 mL/min), D is inner diameter of syringe barrel (8.66 mm), and d is inner diameter of needle (0.337 mm).

### Cryo-SEM imaging

To evaluate depot morphology, PRECISE depots were prepared as described above and incubated under the corresponding in vitro release conditions. At selected time points, depots were collected, gently rinsed to remove residual release medium, and processed for cryo-SEM.

For cryo-SEM, hydrated depots were mounted on cryo-SEM specimen holders and rapidly frozen in liquid nitrogen slush to preserve the hydrated internal morphology. Frozen samples were fractured to expose the depot cross-section, transferred under vacuum to the cryo-preparation chamber, briefly sublimated to remove surface ice, and sputter-coated with platinum/palladium. Samples were maintained below −120 °C during transfer and imaging. Cryo-SEM images were acquired using a field-emission scanning electron microscope equipped with a cryo-stage at an accelerating voltage of 3–5 kV.

### Molecular docking study

To evaluate the binding affinities and interactions between TAC and excipients containing long carbon chains, benzene rings, and heterocyclic rings, molecular docking simulations were performed using PyMOL 2.2.5, AutoDock Vina, and Biovia Discovery Studio 2021 Client. Molecular structures were sourced from the PubChem Database and the Protein Data Bank (RCSB PDB) and converted to .pdb format using PyMOL. These structures were then imported into AutoDock Vina for docking simulations, employing a grid box approach with dimensions of 15 x 15 x 15 Å along the X-, Y-, and Z-axes. Binding affinities, expressed in kcal/mol, were calculated by AutoDock Vina,(*48*) which also generated complex files for each excipient-TAC interaction. Subsequent analyses of these complexes were conducted using Biovia Discovery Studio 2021 Client to identify and characterize specific interactions between TAC and the excipients.

### Pharmacokinetic sampling and drug quantification

For the healthy-rat pharmacokinetic studies, TAC-only or TAC+RAPA formulations were administered as a single subcutaneous injection. Blood was collected from the saphenous on Days 0, 1, 2, 4, 7, 11, 14, 21, and 28; additional samples were collected from TAC+RAPA-treated animals on Days 35, 42, 49, and 56 for TAC quantification into K₂EDTA-coated tubes and analyzed as whole blood.

Tacrolimus concentrations in K₂EDTA whole blood were quantified by liquid chromatography coupled to quadrupole time-of-flight mass spectrometry (LC–QTOF-MS). For sample preparation, 50 µL of whole blood was mixed with 50 µL of 0.1 M ZnSO₄, followed by 200 µL of methanol containing 50 ng/mL ascomycin as an internal standard. Samples were vortexed and centrifuged at 22,000 × g for 10 min, and the resulting supernatants were analyzed.

Chromatographic separation was performed using an Agilent 1290 Infinity UHPLC coupled to an Agilent 6530 Q-TOF mass spectrometer equipped with a Dual AJS electrospray ionization source. Analytes were separated on a Phenomenex Kinetex C18 core-shell column (100 × 2.1 mm, 2.6 µm, 100 Å) maintained at 60 °C. Mobile phase A consisted of 10 mM ammonium formate with 0.1% formic acid in water, and mobile phase B consisted of 10 mM ammonium formate with 0.1% formic acid in acetonitrile. The flow rate was 0.300 mL/min with a 20 µL injection volume. The gradient was as follows: 0–0.5 min, 2% B; 0.5–5.5 min, linear increase to 98% B; 5.5–7.0 min, 98% B; 7.1 min, return to 2% B; and 7.1–10.0 min, re-equilibration at 2% B.

Mass spectrometry was performed in positive electrospray ionization mode using nitrogen as the source gas, with the following parameters: drying gas, 8 L/min at 350 °C; sheath gas, 11 L/min at 400 °C; nebulizer pressure, 45 psi; capillary voltage, 3,500 V; and nozzle voltage, 50 V. Tacrolimus and ascomycin were monitored at m/z 821.5 and 809.5158, respectively.

Rapamycin concentrations in K₂EDTA whole blood were quantified using LC–QTOF-MS/MS. For sample preparation, 100 µL of whole blood was combined with 200 µL of extraction solution containing 10 ng/mL ascomycin and 0.04 M ZnSO₄ in 80:20 methanol:water. Samples were vortexed for 30 s and centrifuged at 14,000 × g for 10 min, and the resulting supernatants were analyzed.

Chromatographic separation was performed using the same Agilent 1290 Infinity UHPLC–Agilent 6530 Q-TOF system and C18 column described above. Mobile phases, flow rate, injection volume, column temperature, and gradient conditions were identical to those used for TAC analysis.

Mass spectrometry was performed in positive electrospray ionization mode with the following parameters: drying gas, 8 L/min at 300 °C; sheath gas, 11 L/min at 350 °C; nebulizer pressure, 45 psi; capillary voltage, 3,500 V; nozzle voltage, 500 V; fragmentor voltage, 130 V; skimmer voltage, 65 V; and octopole RF voltage, 750 V. Data were acquired using targeted MS/MS in positive-ion mode with 2 GHz extended dynamic range. Rapamycin was quantified using the transition m/z 931.58899 [M+NH₄]⁺ → 864.5746, with ascomycin (m/z 809.5158 [M+NH₄]⁺) used as the internal standard.

Calibration curves were linear (R² > 0.999) over concentration ranges of 1.5–200 ng/mL for TAC and 1.5–50 ng/mL for RAPA. The lower limit of quantification was 1.56 ng/mL for both analytes. Data were processed using Agilent MassHunter Quantitative Analysis 10.0, and concentrations were reported as ng/mL whole blood.

### NMP efflux measurement

PRECISE depots were prepared and incubated under the same conditions used for the in vitro release studies. At predetermined time points, release medium was collected and replaced with fresh medium. NMP concentrations in the collected release medium were quantified by reverse-phase HPLC with UV detection at 220 nm using a C18 column (4.6 × 250 mm, 5 μm). Samples (20 μL) were analyzed at room temperature using an isocratic mobile phase consisting of 0.1% (v/v) trifluoroacetic acid in water and acetonitrile (68:32, v/v) at a flow rate of 0.5 mL/min. NMP concentrations were determined from an external calibration curve, and cumulative NMP release was calculated as a percentage of the initial NMP content in each formulation. Experiments were performed in triplicate.

### In vivo depot degradation by IVIS imaging

To evaluate in vivo depot degradation, TAC+RAPA PRECISE depots were prepared using Cy7- conjugated PLGA (85:15 lactide:glycolide, 10–15 kDa). The polymer fraction consisted of 84.8% Cy7- conjugated PLGA and 15.2% unconjugated PLGA. A total of 223.5 mg PLGA was dissolved in 268.4 µL N-methyl-2-pyrrolidone (NMP) and mixed until a homogeneous solution was obtained. TAC and RAPA were incorporated at the same concentrations used for the TAC+RAPA PRECISE formulation (28 mg/mL TAC and 24 mg/mL RAPA, respectively). Formulations were allowed to cool to room temperature, loaded into 1-mL syringes fitted with 23G needles, and approximately 100 µL was injected subcutaneously into each rat (n = 3).

Depot-associated Cy7 fluorescence was monitored noninvasively using an IVIS imaging system at predetermined time points following injection. For each animal, a region of interest was drawn over the depot site, and fluorescence signal was quantified as total radiant efficiency. Signal at each time point was normalized to the initial post-injection signal for the same animal to track loss of depot- associated fluorescence over time.

### Depot retrieval

Healthy male Lewis rats received a single subcutaneous PRECISE injection (0.25 mL; 7 mg TAC, 6 mg RAPA, n = 3). After a week, the animals underwent surgical depot retrieval on POD 7.

Rats were anesthetized with isoflurane (3–4% induction, 1.5–2% maintenance) and given pre-emptive analgesia (buprenorphine SR, 1.0 mg/kg SC). The injection site was shaved, prepped with alternating povidone-iodine and 70% ethanol, and draped. A 1–1.5 cm skin incision was made directly over the palpable depot. The depot, which had solidified by phase inversion, was exposed by blunt and limited sharp dissection through subcutaneous tissue. The intact implant was freed from the fibrous pocket using fine scissors, avoiding fragmentation, and removed en bloc. Skin was closed with 4-0 nylon interrupted sutures and covered with topical antibiotic ointment. Rats recovered in a warmed cage and received postoperative analgesia (buprenorphine 0.05 mg/kg SC every 12 h for 48 h). No antibiotics were used routinely.

### Rat hindlimb allotransplantation

Orthotopic hindlimb transplants were performed from Brown Norway (BN) donors to Lewis recipients under inhalational anesthesia (SomnoSuite®, Kent Scientific). All procedures were performed under sterile conditions by two microsurgeons working in parallel. For donor procurement, a circumferential skin incision was made proximal to the mid-thigh, iliac vessels and nerves were dissected, muscle groups were divided proximally, and the femur was transected at mid-shaft. The limb was flushed with heparinized saline solution and stored on ice until implantation.

For recipient preparation, the femur was stabilized with an intramedullary pin, and donor iliac vessels were anastomosed end-to-end to the recipient femoral vessels using 10-0 nylon under an operating microscope. The sciatic and femoral nerves were coapted with an epineural technique, musculature was reapproximated with 5-0 Vicryl sutures, and the skin was closed using 5-0 Monocryl. Sustained-release buprenorphine (0.05 mg/kg, SC) was administered pre-operatively.

Rats were randomized into three cohorts: untreated controls (n = 12), daily IP TAC comparator (1 mg/kg/day IP; n = 6), and PRECISE-treated animals (n = 12). PRECISE recipients received a 0.25-mL subcutaneous depot of TAC/RAPA formulation at the time of surgery, divided between dorsal and ventral aspects of the graft, followed by identical monthly maintenance depots from postoperative day (POD) 30 to POD 300 (11 total administrations). Daily IP TAC was administered beginning on POD 0.

Animals were examined daily for clinical signs of rejection, including edema, erythema, eschar formation, and necrosis. Peeling of the skin with light pressure was considered confirmation of Banff grade 3 rejection. Body weight was recorded daily for 2 months and weekly thereafter. Animals were euthanized upon reaching Banff grade 3 rejection or at the study endpoint.

### Porcine gracilis free flap allotransplantation

Domestic crossbred Landrace × Yorkshire × Duroc (L × Y × D) pigs (6–8 weeks old; mean body weight approximately 25 kg) underwent reciprocal heterotopic gracilis myocutaneous free-flap transplantation. Flaps (∼5 × 10 × 4 cm) were transferred to the lateral neck. Donor–recipient pairs were selected to ensure full mismatch at major swine leukocyte antigen (SLA) class I and II loci.

Under general anesthesia, the flap pedicle was anastomosed end-to-end to the recipient carotid artery and jugular vein using 8-0 nylon. Local analgesia (0.25% bupivacaine) was provided at incision sites.

Three treatment cohorts were studied: (i) no immunosuppression; (ii) systemic steroid induction only (intraoperative Solu-Medrol 500 mg IV followed by 400, 300, 200, and 100 mg on POD 1, 2, 3, and 4, respectively; and (iii) PRECISE plus systemic steroid induction. In the PRECISE cohort, TAC/RAPA PRECISE was administered at transplantation and on POD 10, 21, 30, 50, and 70. At transplantation, six 0.25-mL injections (total 1.5 mL) delivered 42 mg TAC and 36 mg RAPA per animal. Maintenance administrations consisted of four 0.25-mL injections (total 1.0 mL), delivering 28 mg TAC and 24 mg RAPA per animal.

Postoperative assessments were performed under Telazol sedation (5–8 mg/kg IM) on POD 1, 2, 7, 14, and weekly thereafter. Animals were euthanized under isoflurane overdose at study endpoints or upon reaching Banff grade 3 rejection. At each assessment, grafts were evaluated for perfusion, color, temperature, edema, and skin integrity.

### Swine leukocyte antigen (SLA) genotyping and donor–recipient matching

All animals underwent SLA genotyping prior to selection as previously described (PMID 19392823 and 20121817) with modifications made to the typing primer panels to broaden allele coverage and enhance typing resolution in outbred farm pigs. The class I genes SLA-1, SLA-2, and SLA-3, and the class II genes DRB1, DQB1, and DQA were analyzed. High-resolution, allelic SLA typing was inferred based on previous literature and reports [PMID 15713212, 16305679, 19317739]. Donor-recipient pairs were blood-type compatible and fully mismatched for their SLA class I and class II antigens.

### Histopathology

Skin punch biopsies were obtained at defined time points and at necropsy (2 mm in rats; 6 mm in pigs). Specimens were fixed in 10% neutral-buffered formalin, paraffin-embedded, sectioned (5 µm), and stained with hematoxylin–eosin (H&E) for routine histopathologic evaluation. Additional samples were snap-frozen in OCT for IHC.

Rejection severity was graded according to the modified Banff VCA criteria(*28*), defined as follows: Grade 0, no or rare infiltrates; Grade I, mild, focal infiltrates without epidermal necrosis; Grade II, moderate, multifocal infiltrates with early epidermal necrosis; Grade III, severe, diffuse infiltration with epidermal necrosis; and Grade IV, necrotizing acute rejection. Chronic rejection was assessed by quantifying intimal hyperplasia (IH) and luminal occlusion on H&E- and elastin-stained vessels using ImageJ software.

### Flow cytometry for immune cell phenotyping

For immunophenotyping, PBMCs were isolated from postoperative samples using Ficoll-Hypaque (density 1.077 g/L; TBD Sciences). Cells were washed and resuspended in flow cytometry staining buffer, then stained with fluorescently labeled anti-rat antibodies targeting cell surface markers CD4, CD8, CD25, and the intracellular transcription factor FoxP3 (BD Biosciences, San Diego, CA) for 30 min at 2–8 °C protected from light. After surface staining, cells were washed, fixed, and permeabilized using a transcription-factor fixation/permeabilization buffer set for 40–50 min at 2–8 °C protected from light. Cells were then washed with permeabilization/wash buffer and stained intracellularly with anti-FoxP3 antibody in permeabilization/wash buffer for 40–50 min at 2–8 °C protected from light. Compensation was performed using compensation beads. Stained cells were acquired using a BD LSR Fortessa cytometer, and the data were analyzed with FlowJo software (Tree Star, Ashland, OR). Lymphocytes were first identified by forward- and side-scatter profiles, followed by singlet gating. Treg populations were quantified using a sequential gating strategy based on CD4⁺ cells and CD25⁺FoxP3⁺ expression. The same gating strategy was applied consistently across all samples within each analysis.

### Mixed lymphocyte reaction (MLR) assay in rats

PBMCs from PRECISE–treated long-term surviving Lewis recipients and naïve Lewis rats were isolated by Ficoll-Hypaque density gradient and used as responders. Responders were labeled with CFSE (Thermo Fisher Scientific) at 5 μM for 10 minutes at room temperature, quenched by adding five volumes of complete medium, washed twice, and resuspended in complete RPMI-1640. Donor- strain BN splenocytes were prepared by mechanical dissociation, subjected to ACK red blood cell lysis, washed, and irradiated at 3,000 rad to serve as stimulators. Responder and stimulator cells were combined at a 2:1 ratio in 96-well U-bottom plates in RPMI-1640 media with 10% fetal bovine serum, 1% penicillin–streptomycin, 2 mM L-glutamine, and 50 μM 2-mercaptoethanol. Cultures were incubated for 7 days at 37 °C in 5% CO₂. At the end of the culture period, proliferation was quantified by CFSE dilution on a BD LSR Fortessa cytometer and analyzed in FlowJo.

### Hematology, serum chemistry, and cytokine profiling

Blood samples were collected longitudinally during the rat and porcine VCA studies for complete blood count (CBC), serum chemistry, and cytokine analyses. In the rat study, samples from untreated animals were collected on POD 0–6. Samples from daily IP TAC- and PRECISE-treated animals were collected on POD 0, 7, 30, 60, 90, 120, 150, 180, 210, 240, 270, and 300.

CBC analysis included WBC, neutrophils, lymphocytes, monocytes, eosinophils, RBC, hemoglobin, hematocrit, platelets, MCV, MCHC, and MPV. Serum chemistry analysis included creatinine, BUN, ALT, AST, ALP, total bilirubin, total protein, albumin, glucose, cholesterol, triglycerides, sodium, potassium, calcium, and phosphorus. Circulating cytokines were quantified using a rat-specific multiplex bead assay on a Luminex MAGPIX platform (Luminex, Austin, TX) according to the manufacturer’s instructions. The rat cytokine panel included IL-2, TNF-α, IL-1β, IL-6, IL-4, IFN-γ, IL-17A, IL-10, MCP-1, fractalkine, GM-CSF, LIX/CXCL5, CINC-1/IL-8, and B7-2.

For the porcine study, blood samples were collected from the auricular vein. Samples from untreated animals were collected on POD 0–4, and samples from the steroid-only group were collected on POD 0–6. Samples from PRECISE-treated animals were collected on POD 0, 1, 2, 3, 4, 10, 20, 30, 50, 70, 80, and 90.

CBC analysis included WBC, neutrophils, lymphocytes, monocytes, eosinophils, hemoglobin, hematocrit, platelets, MCV, and MCHC. Serum chemistry analysis included creatinine, BUN, ALT, AST, ALP, GGT, total bilirubin, total protein, albumin, glucose, sodium, potassium, cholesterol, triglycerides, calcium, and phosphorus. Porcine cytokines were quantified using a MILLIPLEX MAP Porcine Cytokine/Chemokine Magnetic Bead Panel (MilliporeSigma) on a Luminex MAGPIX platform according to the manufacturer’s instructions. The multiplex panel included IL-2, TNF-α, IL-1β, IL-6, IL-4, IFN-γ, IL-8/CXCL8, IL-10, IL-12, and IL-18. The standard MILLIPLEX porcine panel specifically contains these analytes. Merck Millipore IL-17A was quantified using a porcine-specific ELISA, and CRP was quantified using a MILLIPLEX Porcine Acute Phase Protein Panel (PAPP1-160K, MilliporeSigma). The latter is a Luminex-based porcine assay specifically validated for CRP and haptoglobin.

## Acknowledgement

We thank the veterinary and animal care staff at Brigham and Women’s Hospital and Wake Forest University for their support with rat and porcine studies and postoperative monitoring. We also acknowledge the use of ChatGPT (OpenAI) for language refinement and assistance in improving the clarity and readability of this manuscript. **Funding:** This work was supported by the National Institutes of Health (NIH) grant R21DA057701 (to N.J.); the Department of Defense Award W81XWH-15-2-0047 (to J.M.K. and V.G.), by the Department of Anesthesiology, Perioperative, and Pain Medicine at Brigham and Women’s Hospital (to N.J. and J.M.K.); by the Nano & Material Technology Development Program through the National Research Foundation of Korea (NRF) funded by the Ministry of Science and ICT (RS-2024-00405574) (to S.L.); by the Harvard University Center for AIDS Research (CFAR) (to S.L.); and by the NIH/NIAID K99/R00 Pathway to Independence Award K99AI191990 (to S.L.). This research was, in part, funded by the Advanced Research Projects Agency for Health (ARPA-H; OTA #1AY2AX000056). The views and conclusions contained in this document are those of the authors and should not be interpreted as representing the official policies, either expressed or implied, of the United States Government. **Author contributions:** Conceptualization: S.L., J.J., N.J., J.M.K., V.G. Methodology: S.L., F.Z., Y.K., J.J., D.P., G.J., S.V., W.Z., N.J., V.G. Software: D.P. Investigation: S.L., F.Z., Y.K., J.J., H.K., S.B., G.M., D.P., J.N.L., L.Z., A.W., K.C., N.L., P.S., E.A., H.M.B., O.S., X.C., K.S., M.D., E.S., A.G., G.J., S.V., W.Z., V.G. Formal analysis: S.L., J.J., D.P., W.Z. Data curation: S.L., N.J. Visualization: S.L., F.Z., Y.K., G.M. Writing – original draft: S.L. Writing – review and editing: S.L., F.Z., J.M.K., N.J., V.G. Supervision: J.M.K., N.J., V.G. Project administration: S.L., J.M.K., N.J., V.G. Funding acquisition: S.L., J.M.K., N.J. **Competing interests:** S.L., J.J., J.M.K. N.J., and V.G. are inventors on a pending patent based on the PRECISE formulation described in this manuscript. J.M.K. has served as a paid consultant and/or holds equity in multiple biotechnology companies, including those listed at https://www.karplab.net/team/jeff-karp (for example, Stempeutics, Sanofi, Celltex, LifeVaultBio, Takeda, Ligandal, Camden Partners, Stemgent, Biogen, Pancryos, Element Biosciences, Frequency Therapeutics, Corner Therapeutics, Quthero, and Mesoblast). These interests have been reviewed and are managed in accordance with institutional conflict-of-interest policies. **Data and materials availability:** All data needed to evaluate the conclusions of this study are available in the main text and/or the Supplementary Materials. Additional data, analytical code, and materials, including PRECISE formulations and related protocols, are available from the corresponding authors upon reasonable request and completion of a material transfer agreement, where applicable.

