## Supplementary material for "A Precisely Controlled Long-Acting Immunosuppression Platform Enables Prolonged Survival of Vascularized Composite Allografts": Supplemetary data

**This PDF file includes:**

**Supplementary Figure 1 to 16**

**Supplementary Table 1**

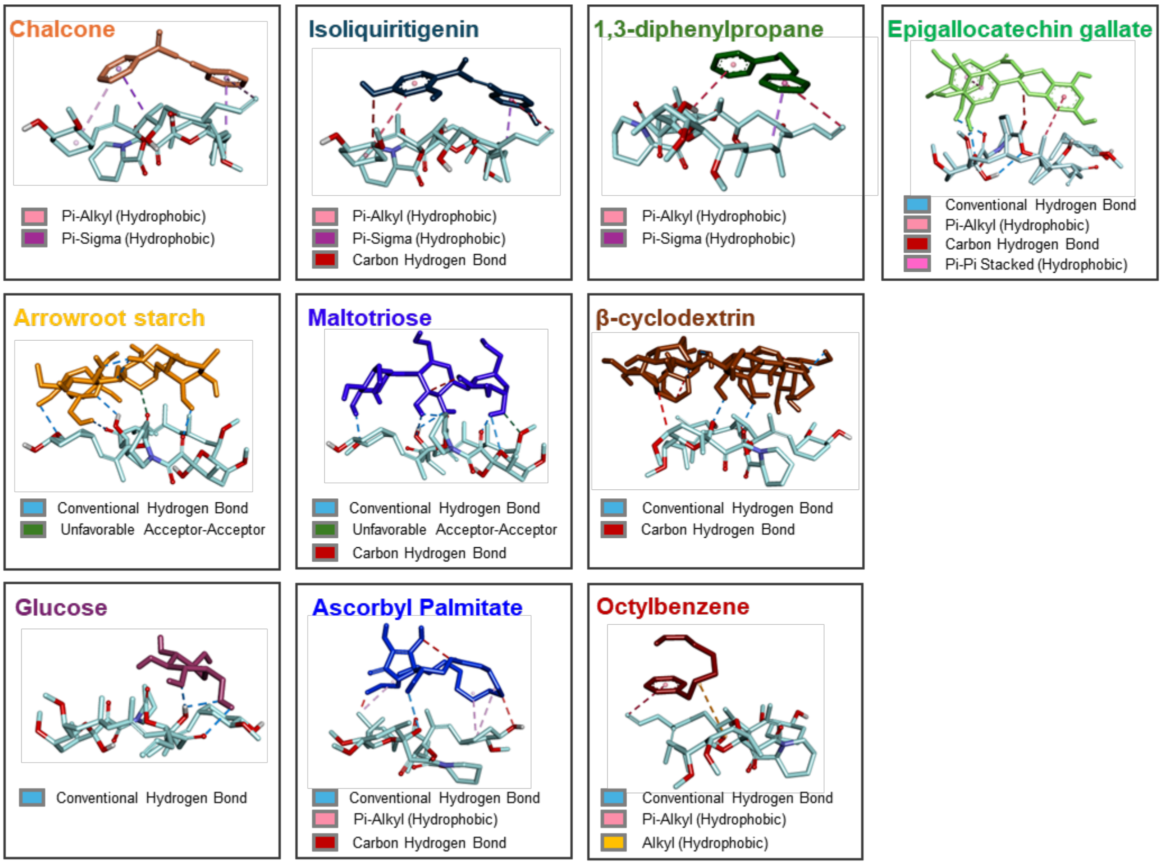


**Figure S1. Representative *in silico* docking poses (AutoDock Vina) of TAC with GRAS-listed IMMs.** Compounds were classified into three structural groups—aromatic polyphenols, carbohydrates, and long-chain hydrophobes—for experimental validation.

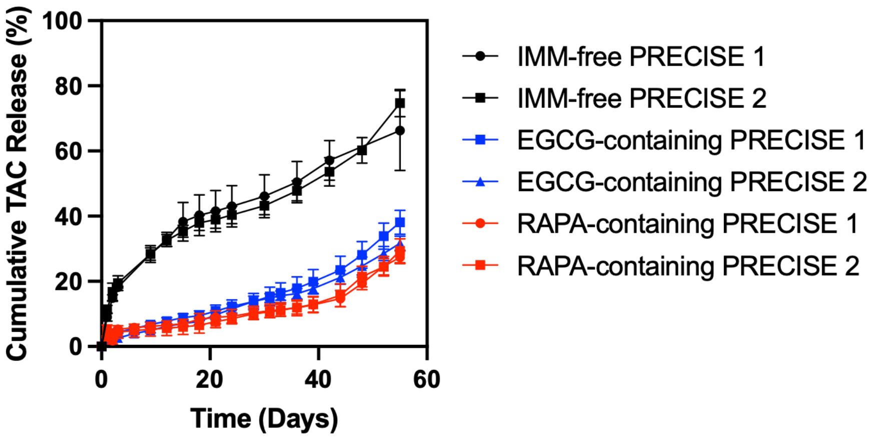


**Figure S2. Reproducible TAC release from representative PRECISE formulations.** In vitro release studies confirmed consistent TAC release profiles across PRECISE formulations without IMM, with EGCG, and with RAPA. Data are presented as mean ± standard deviation (n = 3 samples per formulation).

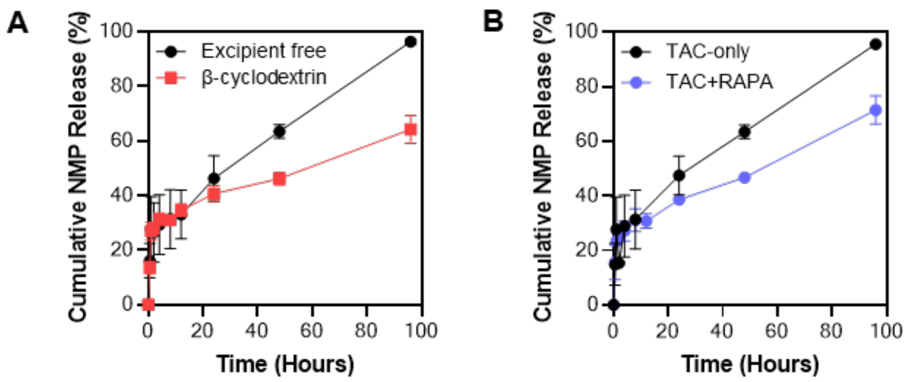


**Figure S3. Drug–excipient modulation of NMP efflux during depot formation. (A)** Cumulative NMP release from PRECISE depots with or without β-cyclodextrin. **(B)** Cumulative NMP release from TAC-only and TAC+RAPA PRECISE depots showing that RAPA co-loading attenuates NMP efflux, suggesting that TAC-RAPA interactions modulate solvent dynamics during phase inversion. Data are presented as mean ± standard deviation (n = 3 samples per formulation).

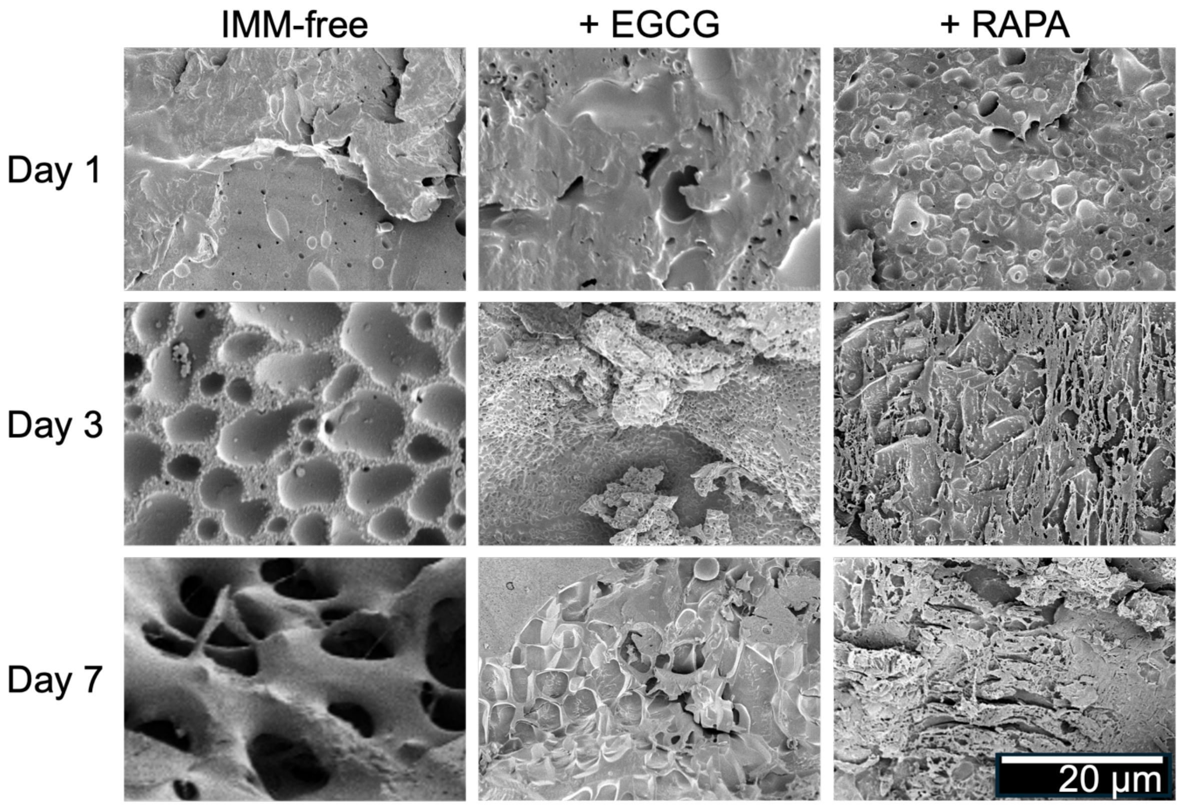


**Figure S4. Cryo-SEM characterization of depot microstructure over time.** Representative cryo-SEM images of TAC-loaded IMM-free PLGA/NMP control, EGCG-containing PRECISE depots, and RAPA-containing PRECISE depots at days 1, 3, and 7, showing time- and formulation-dependent differences in depot microstructure and pore morphology. Representative images were obtained from n = 3 depots per formulation at each time point.

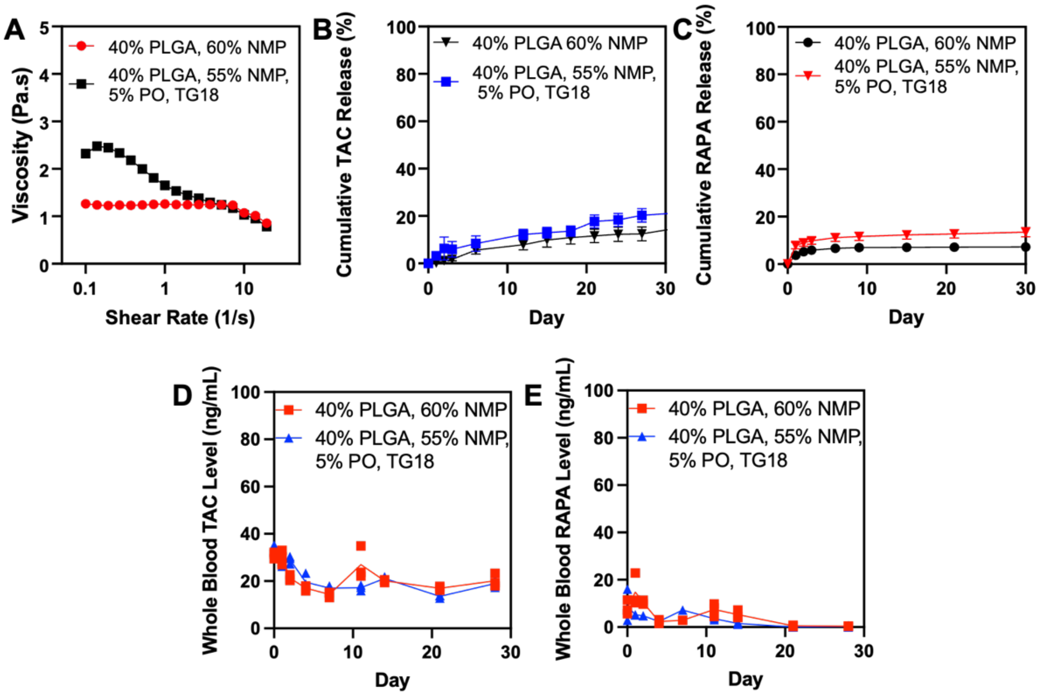


**Figure S5. Shear-thinning formulation for on-bed dosing preserves PRECISE release kinetics both in vitro and in vivo. (A)** Rotational rheometry of TAC+RAPA PRECISE with and without palm oil/TG-18, showing increased viscosity and shear-thinning behavior following incorporation of palm oil/TG-18. **(B–C)** In vitro cumulative release of **(B)** TAC and **(C)** RAPA from TAC+RAPA PRECISE with and without palm oil/TG-18. Data are shown as mean ± SEM, n=3. **(D–E)** Whole-blood pharmacokinetic profiles following a single subcutaneous injection of TAC+RAPA PRECISE and TAC+RAPA PRECISE containing palm oil/TG-18 in healthy rats, showing comparable **(D)** TAC and **(E)** RAPA exposure between formulations. The assay limit of detection (LOD) was 1.56 ng/mL. For RAPA in the standard TAC+RAPA PRECISE group, all animals remained above the LOD through Day 14 and were below the LOD by Day 21. In the palm oil/TG-18 group, RAPA was below the LOD in 1 of 3 animals on Days 1, 4, and 7; 2 of 3 animals on Days 2 and 14; and all animals by Day 21. Rheology in (A) was performed using n = 3 independent samples per formulation. Pharmacokinetic data in (D-E) represent n = 3 rats per formulation.

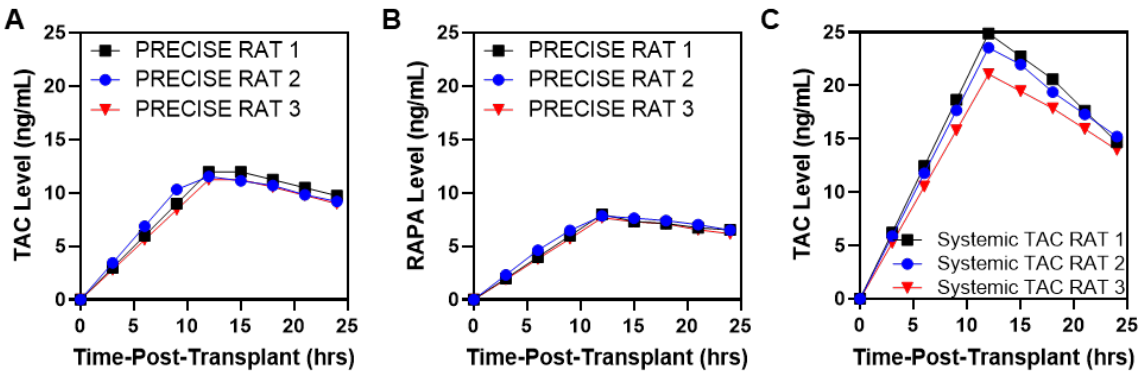


**Figure S6. Early-phase (0–24 h) pharmacokinetics in rats.** Early-phase (0–24 h) pharmacokinetic profiles of **(A)** TAC and **(B)** RAPA after a single subcutaneous PRECISE injection, compared with **(C)** daily intraperitoneal TAC. Concentrations were sampled over 24 h and used for $AUC_{0\text{-}24\,h}$ calculations shown in Figure 4D. Pharmacokinetic profiles represent n = 3 rats per treatment group.

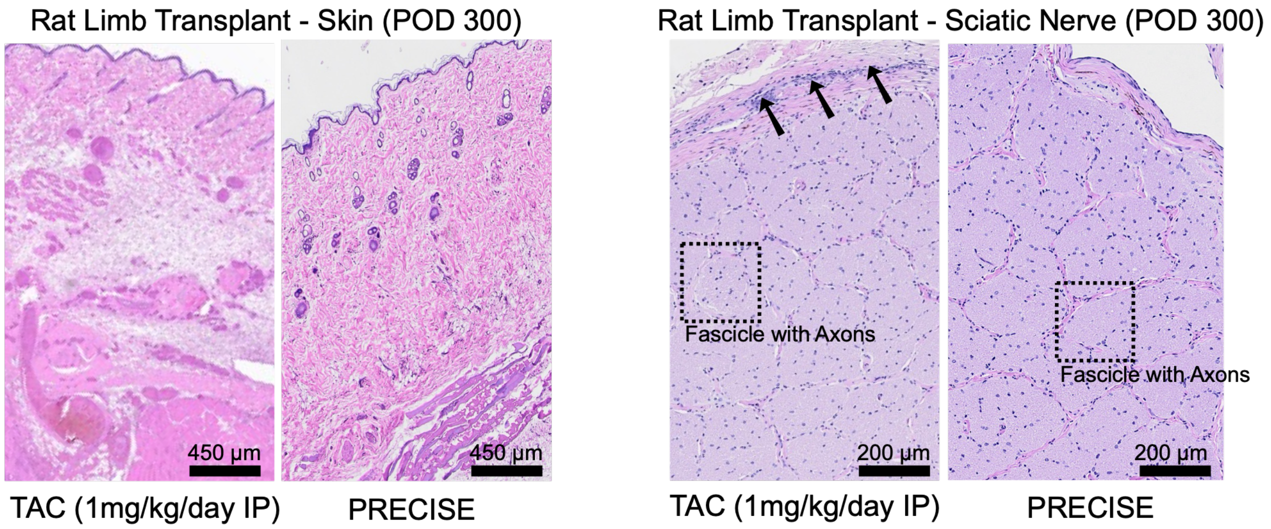


**Figure S7. H&E analysis of transplanted skin and sciatic nerve in daily IP TAC- and PRECISE-treated rat grafts.** Representative H&E-stained sections of transplanted skin and sciatic nerve from daily IP TAC- and PRECISE-treated rat hindlimb allografts at POD 300. Compared with PRECISE-treated grafts, daily IP TAC-treated grafts showed dermal tissue disruption and perineurial/epineurial mononuclear inflammatory infiltrates around nerve fascicles. PRECISE-treated grafts showed preserved dermal architecture and organized nerve fascicles with identifiable axon-containing fascicular regions. Arrows indicate inflammatory infiltrates around nerve fascicles. Dashed boxes indicate representative axon-containing fascicular regions. Scale bars: 450 µm for skin and 200 µm for sciatic nerve.

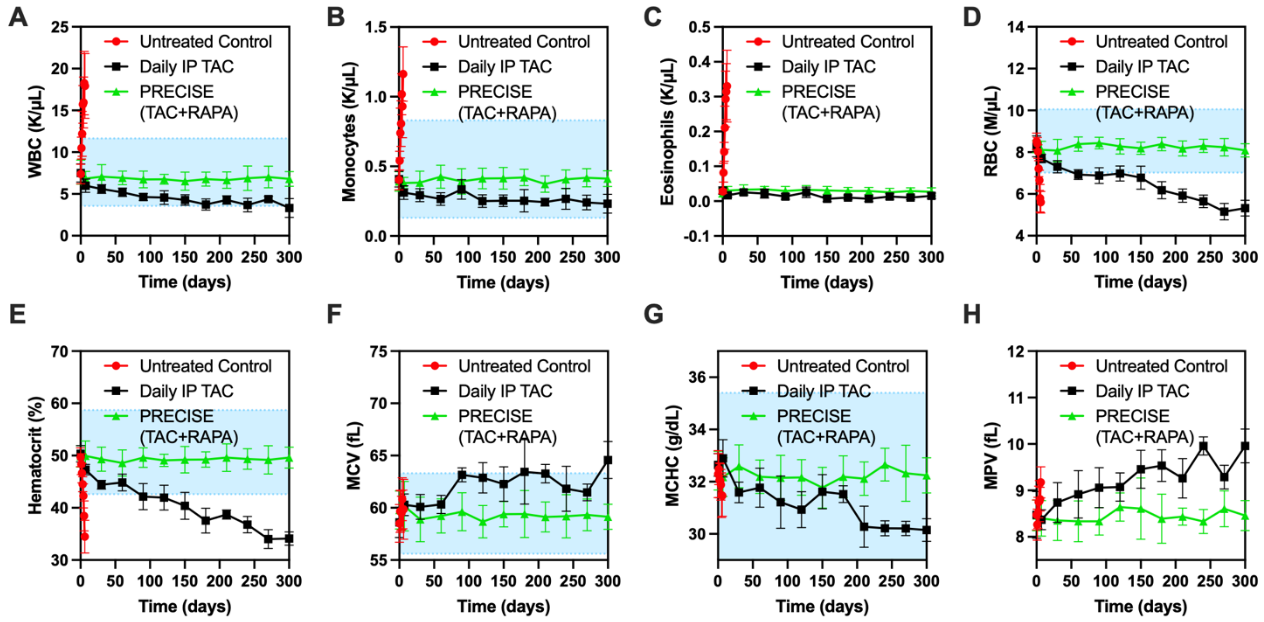


**Figure S8. PRECISE maintains additional hematologic parameters within or near reported reference ranges in the rat VCA model.** Additional CBC parameters included **(A)** WBC, **(B)** monocytes, **(C)** eosinophils, **(D)** RBC, **(E)** hematocrit, **(F)** MCV, **(G)** MCHC, and **(H)** MPV. Daily IP TAC-treated animals showed decreases in WBC, RBC, hematocrit, and MCHC at later time points, whereas these parameters generally remained within or near reported rat reference ranges in PRECISE-treated animals. Blue shaded regions indicate rat reference ranges where available *(50)*. Group sizes were n=12 for untreated controls, n=6 for daily IP TAC-treated animals, and n=12 for PRECISE-treated animals.

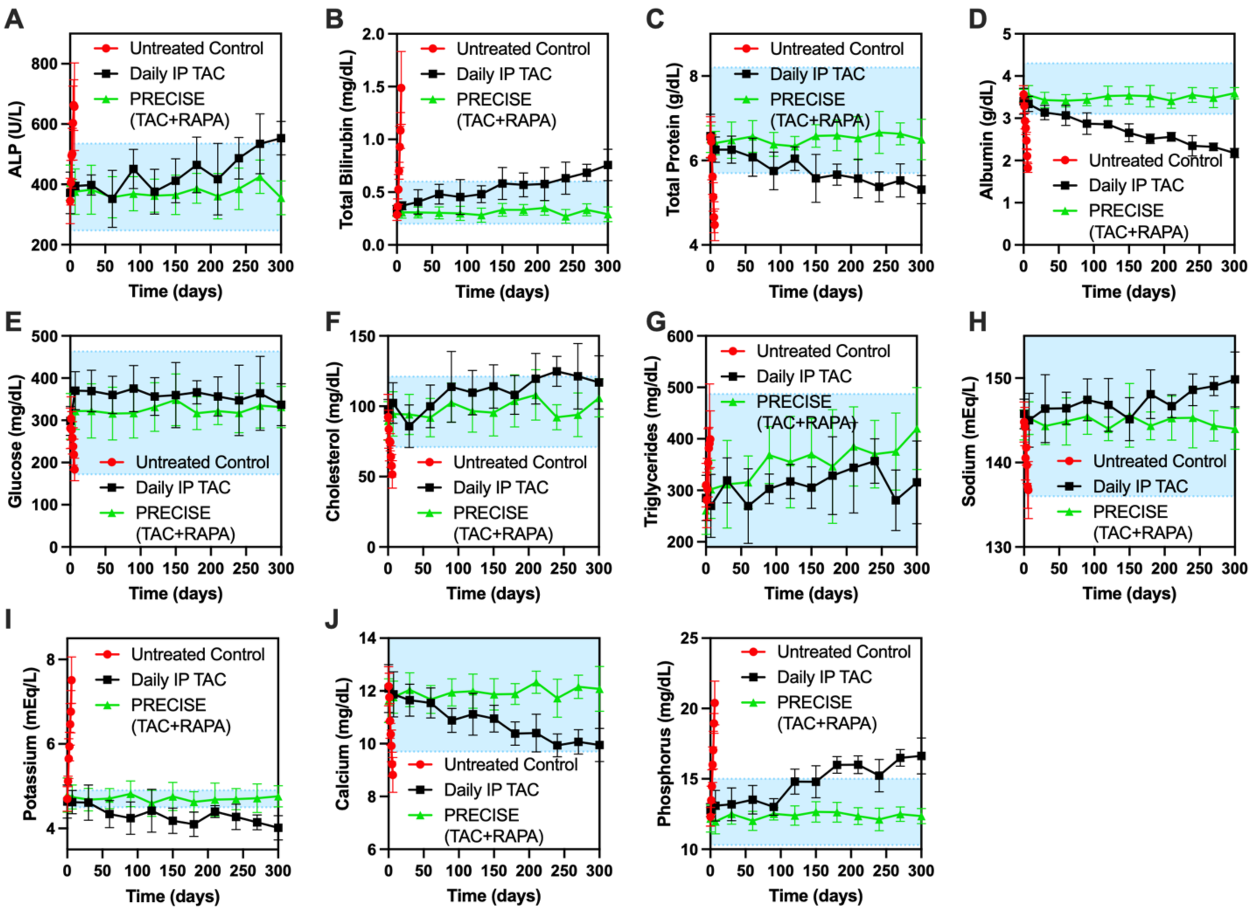


**Figure S9. PRECISE maintains additional serum chemistry parameters within or near reported reference ranges in the rat VCA model.** Additional serum chemistry parameters included **(A)** ALP, **(B)** total bilirubin, **(C)** total protein, **(D)** albumin, **(E)** glucose, **(F)** cholesterol, **(G)** triglycerides, **(H)** sodium, **(I)** potassium, **(J)** calcium, and **(K)** phosphorus. Daily IP TAC-treated animals showed increases in ALP, total bilirubin, and phosphorus and decreases in total protein, albumin, potassium, and calcium at later time points, whereas these parameters generally remained within or near reported rat reference ranges in PRECISE-treated animals. Blue shaded regions indicate rat reference ranges where available *(50)*. Group sizes were n=12 for untreated controls, n=6 for daily IP TAC-treated animals, and n=12 for PRECISE-treated animals.

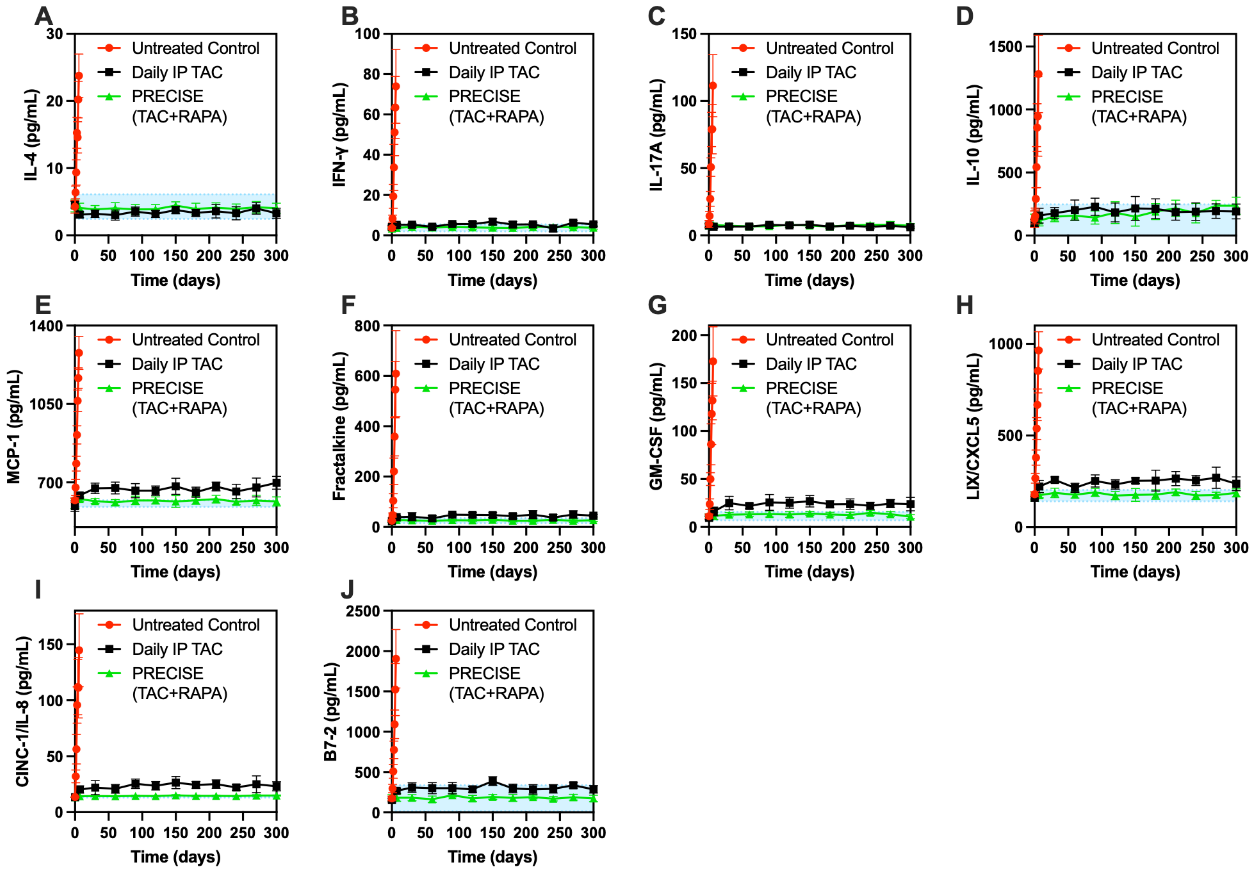


**Figure S10. PRECISE maintains additional circulating cytokines within or near reported baseline ranges in the rat VCA model.** Additional cytokines included **(A)** IL-4, **(B)** IFN-γ, **(C)** IL-17A, **(D)** IL-10, **(E)** MCP-1, **(F)** fractalkine, **(G)** GM-CSF, **(H)** LIX/CXCL5, **(I)** CINC-1/IL-8, and **(J)** B7-2. Cytokine concentrations in PRECISE-treated animals generally remained within or near reported rat baseline ranges throughout the study. Blue shaded regions indicate published rat baseline ranges where available (*49*). Group sizes were n=12 for untreated controls, n=6 for daily IP TAC-treated animals, and n=12 for PRECISE-treated animals.

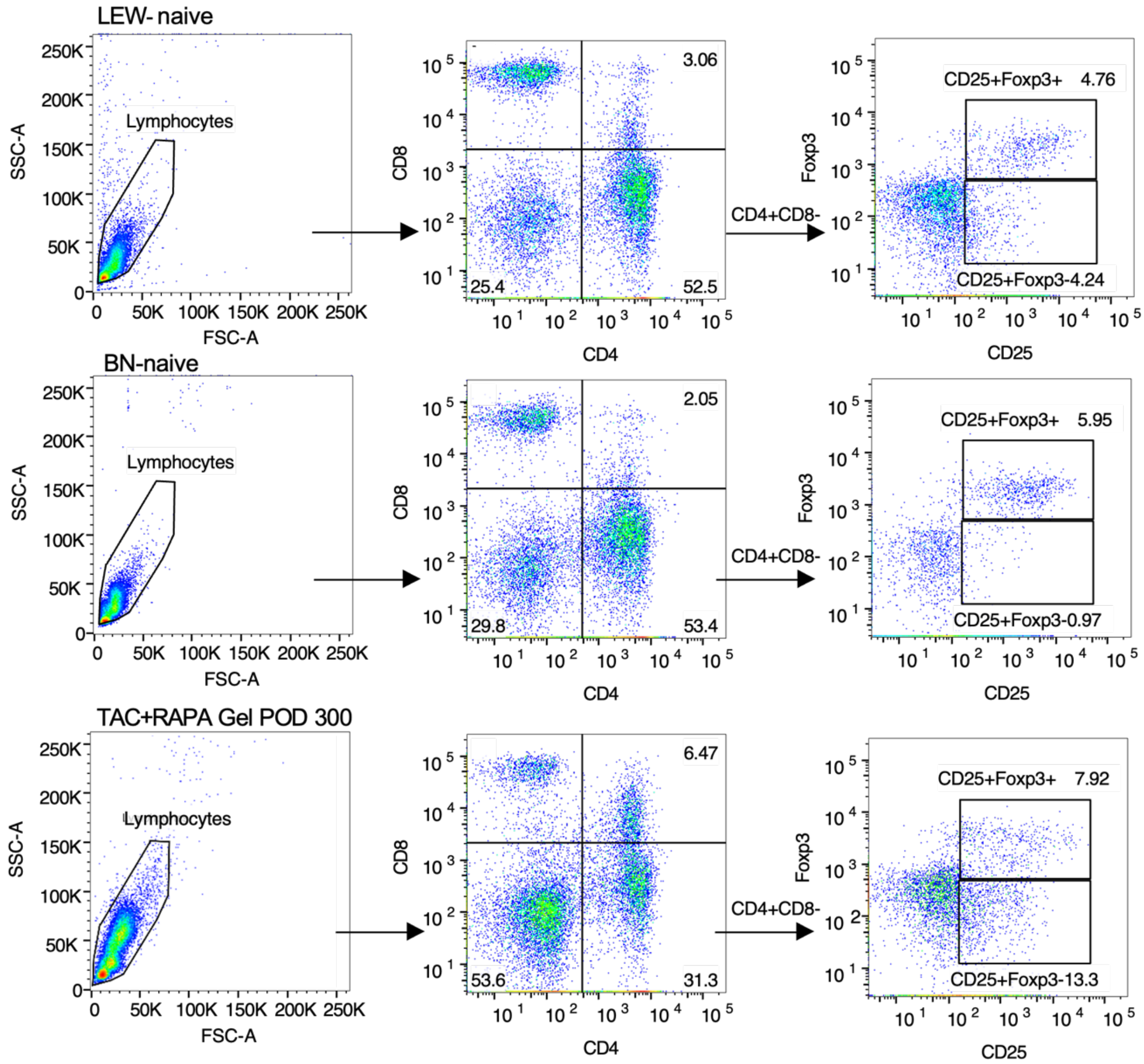


**Figure S11. Flow cytometry gating and longitudinal regulatory T-cell analysis in rats.** Sequential gating strategy for identification of $CD4^{+}CD25^{+}Foxp3^{+}$ Tregs from peripheral blood mononuclear cells (PBMCs). Representative dot plots from naïve (Lewis and Brown Norway) and PRECISE-treated rats at POD 300 showing expansion of the Treg population.

| | | DONOR | | | | | | |
|---|---|---|---|---|---|---|---|---|
| | Animal ID | P21063 A 32.1/62.12a | P21064 A 32.1/32.15b | P21065 A 31.13a/32.1 | P21066 Non-A 40.11/58.WF1 | P21067 Non-A 34.23/40.11 | P21068 Non-A 45.11/59.27 | P21069 A 32.1/58.WF1 |
| RECIPIENT | P21063 A 32.1/62.12a | – | 1 Class II haplotype mismatch | 1 haplotype mismatch | 2 haplotype mismatch | 2 haplotype mismatch | 2 haplotype mismatch | 2 haplotype mismatch |
| | P21064 A 32.1/32.15b | 1 haplotype mismatch | – | 1 haplotype mismatch | 2 haplotype mismatch | 2 haplotype mismatch | 2 haplotype mismatch | 2 haplotype mismatch |
| | P21065 A 31.13a/32.1 | 1 haplotype mismatch | 1 Class II haplotype mismatch | – | 2 haplotype mismatch | 2 haplotype mismatch | 2 haplotype mismatch | 2 haplotype mismatch |
| | P21066 Non-A 40.11/58.WF1 | Blood type incompatible | Blood type incompatible | Blood type incompatible | – | 1 haplotype mismatch | 2 Class I and II haplotype mismatch | Blood type incompatible |
| | P21067 Non-A 34.23/40.11 | Blood type incompatible | Blood type incompatible | Blood type incompatible | 1 haplotype mismatch | – | 2 Class I and II haplotype mismatch | Blood type incompatible |
| | P21068 Non-A 45.11/59.27 | Blood type incompatible | Blood type incompatible | Blood type incompatible | 2 Class I and II haplotype mismatch | 2 Class I and II haplotype mismatch | – | Blood type incompatible |
| | P21069 A 32.1/58.WF1 | 1 haplotype mismatch | 1 Class II haplotype mismatch | 1 haplotype mismatch | 1 haplotype mismatch | 2 haplotype mismatch | 2 haplotype mismatch | – |

**Figure S12. Donor–recipient pairs were selected to maximize immunological disparity using a compatibility matrix based on swine leukocyte antigen (SLA) haplotypes.** SLA Class I (SLA-1, SLA-2, SLA-3) and Class II (DRB1, DQB1) typing was performed using low-resolution PCR and high-resolution sequence-based genotyping. Haplotypes were assigned according to IPD-MHC SLA database nomenclature. SLA-matched pairs and ABO-incompatible pairs were excluded to avoid immunological permissiveness or hyperacute rejection, respectively. Final pairs reflected full Class I and II mismatches. Two reciprocal donor–recipient combinations were selected to assess directionality under maximal allogeneic pressure.

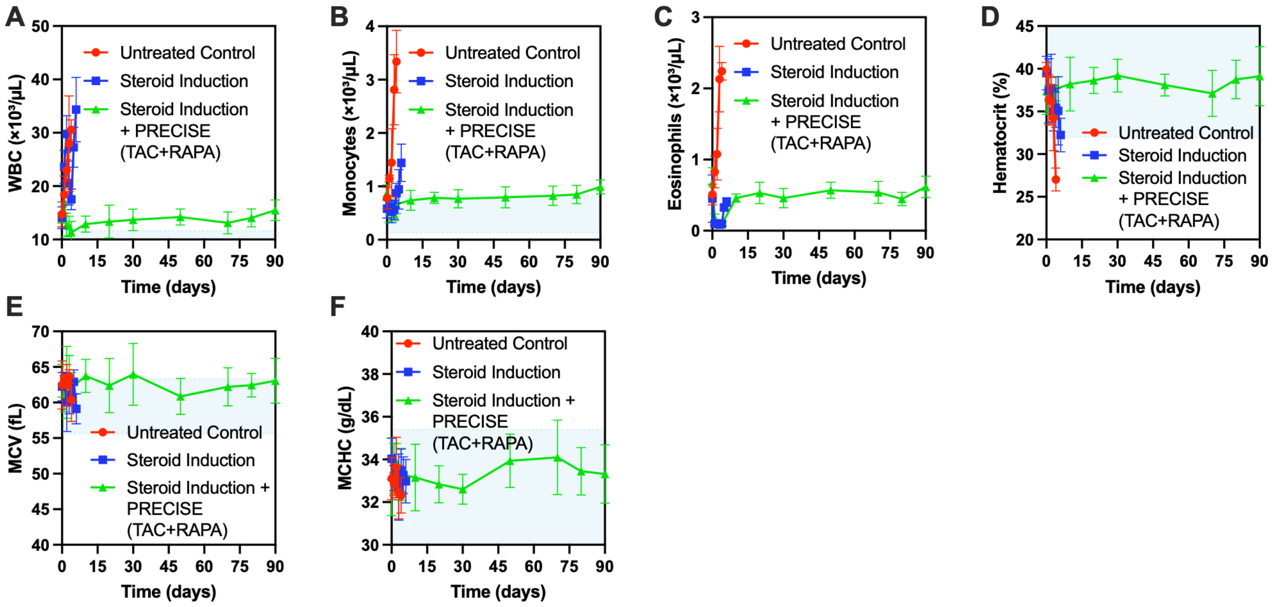


**Figure S13. PRECISE maintains additional hematologic parameters within or near reported reference ranges in the porcine VCA model.** Additional CBC parameters included **(A)** WBC, **(B)** monocytes, **(C)** eosinophils, **(D)** hematocrit, **(E)** MCV, and **(F)** MCHC. These parameters generally remained within or near reported porcine reference ranges in PRECISE-treated animals throughout the study. Blue shaded regions indicate porcine reference ranges where available (*51*). Group sizes were n=4 for untreated controls, n=4 for steroid-induction-only animals, and n=6 for PRECISE + steroid-treated animals.

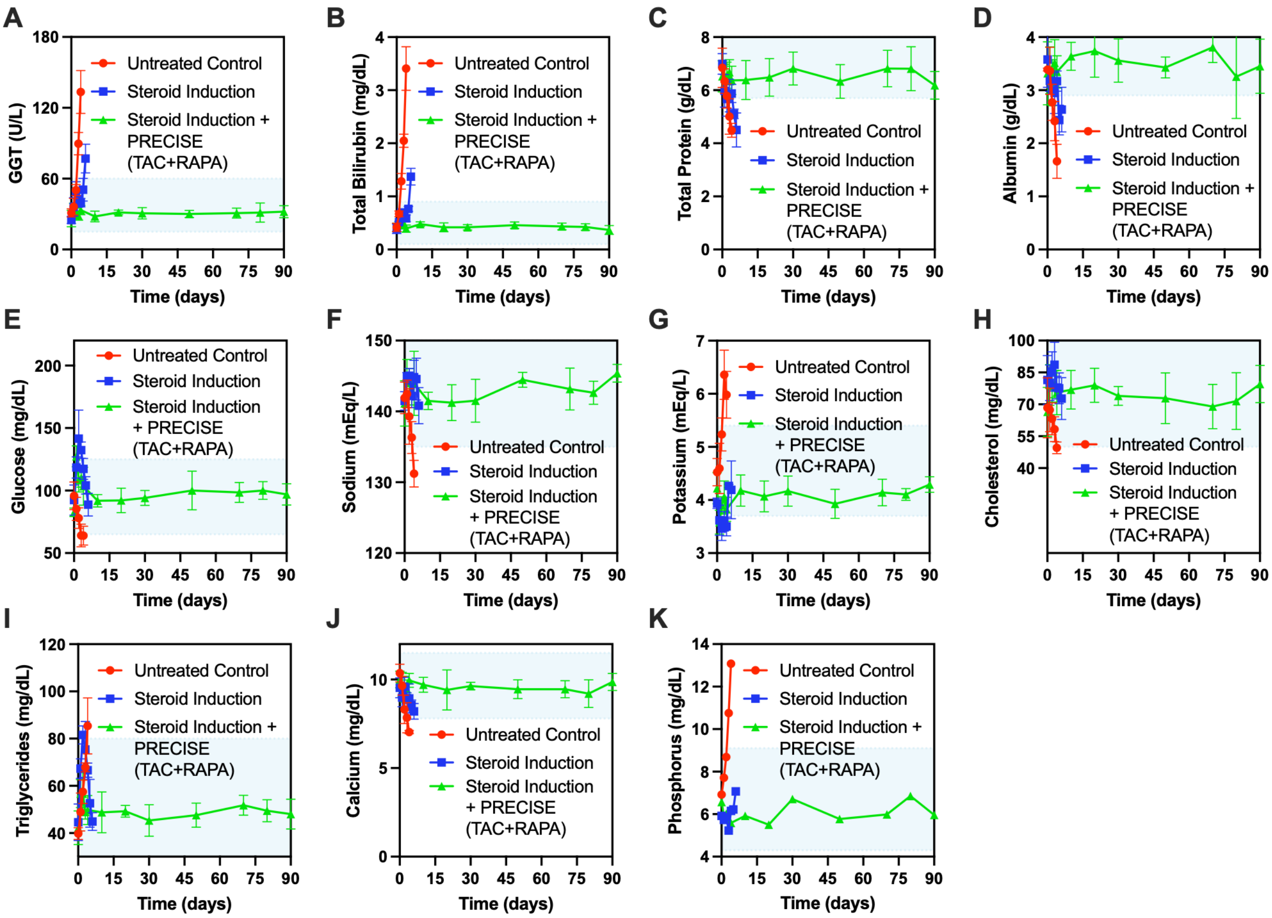


**Figure S14. PRECISE maintains additional serum chemistry parameters within or near reported reference ranges in the porcine VCA model.** Additional serum chemistry parameters included **(A)** GGT, **(B)** total bilirubin, **(C)** total protein, **(D)** albumin, **(E)** glucose, **(F)** sodium, **(G)** potassium, **(H)** cholesterol, **(I)** triglycerides, **(J)** calcium, and **(K)** phosphorus. These parameters generally remained within or near reported porcine reference ranges in PRECISE-treated animals throughout the study. Blue shaded regions indicate porcine reference ranges where available (*51*). Group sizes were n=4 for untreated controls, n=4 for steroid-induction-only animals, and n=6 for PRECISE + steroid-treated animals.

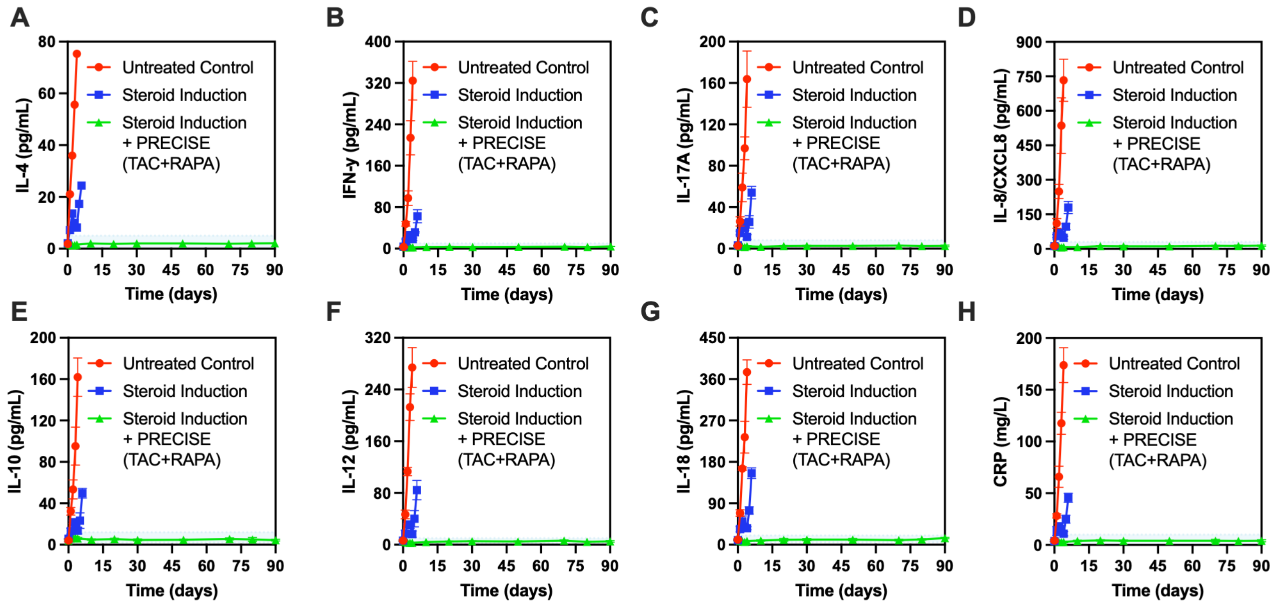


**Figure S15. PRECISE maintains additional circulating cytokine and inflammatory markers within or near reported baseline ranges in the porcine VCA model.** Additional cytokine and inflammatory markers included **(A)** IL-4, **(B)** IFN-γ, **(C)** IL-17A, **(D)** IL-8/CXCL8, **(E)** IL-10, **(F)** IL-12, **(G)** IL-18, and **(H)** CRP. These markers generally remained within or near reported porcine baseline ranges in PRECISE-treated animals throughout the study. Blue shaded regions indicate baseline ranges derived from pre-treatment/control measurements where available. Group sizes were n=4 for untreated controls, n=4 for steroid-induction-only animals, and n=6 for PRECISE + steroid-treated animals.

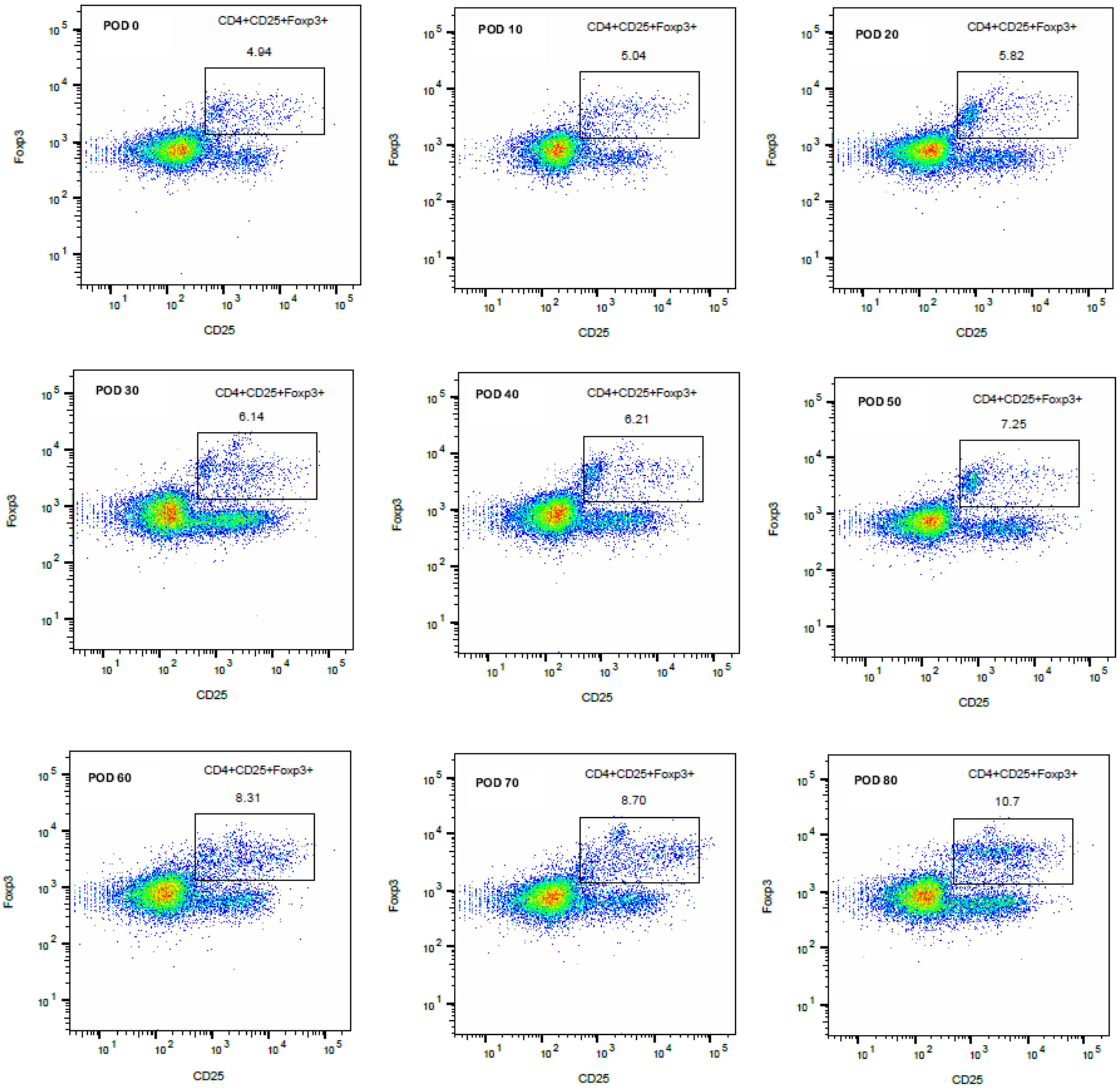


**Figure S16. Flow cytometry of peripheral blood mononuclear cells (PBMCs) in the PRECISE + steroid cohort showed a progressive increase in CD4$^{+}$CD25$^{+}$Foxp3$^{+}$ Tregs from ~5% at baseline (POD 0) to ~11% by POD 80.**

**Table**

**Table S1. HPLC conditions for in-vitro tacrolimus quantification.**

| Time (min) | Methanol (%) | Water (%) | Flow rate (mL/min) |
|---|---|---|---|
| 0 | 70 | 30 | |
| 3 | 70 | 30 | |
| 15 | 90 | 10 | 1 |
| 16 | 90 | 10 | |
| 19 | 70 | 30 | |
